# Asynchronous and coherent dynamics in balanced excitatory-inhibitory populations

**DOI:** 10.1101/2021.08.03.454860

**Authors:** Hongjie Bi, Matteo di Volo, Alessandro Torcini

## Abstract

Dynamic excitatory-inhibitory (E-I) balance is a paradigmatic mechanism invoked to explain the irregular low firing activity observed in the cortex. However, we will show that the E-I balance can be at the origin of other regimes observable in the brain. The analysis is performed by combining extensive simulations of sparse E-I networks composed of *N* spiking neurons with analytical investigations of low dimensional neural mass models. The bifurcation diagrams, derived for the neural mass model, allow to classify the possible asynchronous and coherent behaviours emerging in balanced E-I networks with structural heterogeneity for any finite in-degree *K*. In the limit *N >> K >>* 1 both supra and sub-threshold balanced asynchronous regimes can be observed in our system. Due to the heterogeneity the asynchronous states are characterized by the splitting of the neurons in three groups: silent, fluctuation and mean driven. These features are consistent with experimental observations reported for heterogeneous neural circuits. The coherent rhythms observed in our system can range from periodic and quasi-periodic collective oscillations (COs) to coherent chaos. These rhythms are characterized by regular or irregular temporal fluctuations joined to spatial coherence somehow similar to coherent fluctuations observed in the cortex over multiple spatial scales. The COs can emerge due to two different mechanisms. A first mechanism similar to the pyramidal-interneuron gamma (PING) one, usually invoked for the emergence of *γ*-oscillations. The second mechanism is intimately related to the presence of current fluctuations, which sustain COs characterized by an essentially simultaneous bursting of the two populations. We observe period-doubling cascades involving the PING-like COs finally leading to the appearance of coherent chaos. Fluctuation driven COs are usually observable in our system as quasi-periodic collective motions characterized by two incommensurate frequencies. However, for sufficiently strong current fluctuations we report a novel mechanism of frequency locking among collective rhythms promoted by these intrinsic fluctuations. Our analysis suggest that despite PING-like or fluctuation driven COS are observable for any finite in-degree *K*, in the limit *N >> K >>* 1 these solutions finally result in two coexisting balanced regimes: an asynchronous and a fully synchronized one.

## 1 INTRODUCTION

Cortical neurons are subject to a continuous bombardment from thousands of pre-synaptic neurons, mostly pyramidal ones, evoking post-synaptic potentials of sub-millivolt or millivolt amplitudes (Destexhe and Paré, 1999; Bruno and Sakmann, 2006; Lefort et al., 2009). This stimulation would induce an almost constant depolarization of the neurons leading to a regular firing, However, cortical neurons fire quite irregularly and with low firing rates (Softky and Koch, 1993). This apparent paradox can be solved by introducing the concept of a balanced network, where excitatory and inhibitory synaptic currents are approximately balanced and the neurons are kept near their firing threshold crossing it at random times (Shadlen and Newsome, 1994, 1998). However, the balance should naturally emerge in the network without fine tuning of the parameters and the highly irregular firing observed *in vivo* should be maintained also for large number of connections (in-degree) *K >>* 1. This is possible by considering a sparse excitatory-inhibitory (E-I) neural network composed by *N* neurons and characterized by an average in-degree *K << N* and by synaptic couplings scaling as 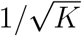 (van Vreeswijk and Sompolinsky, 1996). This scaling as well as many other key predictions of the theory developed in (van Vreeswijk and Sompolinsky, 1996) have been recently confirmed by experimental measurements *in vitro* on a neural culture optogeneitically stimulated (Barral and Reyes, 2016). Furthermore, the authors in (Barral and Reyes, 2016) have shown that the major predictions of the seminal theory (van Vreeswijk and Sompolinsky, 1996) hold also under conditions far from the asymptotic limits where *K* and *N* are large.

The dynamics usually observable in balanced neural networks is asynchronous and characterized by irregular neural firing joined to stationary firing rates (van Vreeswijk and Sompolinsky, 1996; Renart et al., 2010; Litwin-Kumar and Doiron, 2012; Monteforte and Wolf, 2010; Ullner et al., 2020). However, asynchronous regimes characterized by mean driven sub-Poissonian statistics as well as by super-Poissonian one have been reported in balanced homogeneous and heterogeneous networks (Lerchner et al., 2006; Ullner et al., 2020). Furthermore, regular and irregular collective oscillations (COs) have been shown to emerge in balanced networks composed of rate models (van Vreeswijk and Sompolinsky, 1996) as well as of spiking neurons (Brunel, 2000; Ostojic, 2014; Ullner et al., 2018; di Volo and Torcini, 2018; Bi et al., 2020). The balanced asynchronous irregular state has been experimentally observed both *in vivo* and *in vitro* (Shu et al., 2003; Haider et al., 2006) and dynamic balance of excitation and inhibition is observable in the neocortex across all states of the wake-sleep cycle, in both human and monkey (Dehghani et al., 2016). However, this is not the only balanced state observable during spontaneous cortical activity. In particular, balancing of excitation and inhibition appears to be crucial for the emergence of cortical oscillations (Okun and Lampl, 2008; Isaacson and Scanziani, 2011; Le Van Quyen et al., 2016).

In this work we characterize in details the asynchronous regimes and the emergence of COs (population rhythms) in E-I balanced networks with structural heterogeneity. In particular, we consider sparse random networks of quadratic integrate-and-fire (QIF) neurons (Ermentrout and Kopell, 1986) pulse coupled via instantaneous post-synaptic potentials. We compare numerical findings with analytical results obtained in the mean-field (MF) limit by employing an effective low-dimensional neural mass model recently developed for sparse QIF networks (Montbrió et al., 2015; di Volo and Torcini, 2018; Bi et al., 2020).

In the asynchronous regime, our analytical MF predictions are able reproduce the mean membrane potentials and the population firing rates of the structurally heterogeneous network for any finite *K* value. Furthermore, in the limit *N >> K >>* 1 we analytically derive the asymptotic MF values of the population firing rates as well as of the effective input currents. This analysis shows that the system always achieve a balanced dynamics, whose supra or sub-threshold nature is determined by the model parameters. Detailed numerical investigations of the microscopic dynamics allow to identify three different groups of neurons, whose activity is essentially controlled by their in-degrees and by the effective input currents.

In the balanced network we have identified three types of COs depending on the corresponding solution displayed by the neural mass model. The first type, termed O_*P*_ emerges in the MF via a Hopf bifurcation from a stable focus solution. These COs gives rise to collective chaos via a period-doubling sequence of bifurcations. Another type of CO, already reported for purely inhibitory networks (di Volo and Torcini, 2018), denoted as O_*F*_ corresponds in the MF to a stable focus characterized by relaxation oscillations towards the fixed point that in the sparse network become noise sustained oscillations due to fluctuations in the input currents. The last type of COs identified in the finite network are named O_*S*_ and characterized by an abnormally synchronized dynamics among the neurons, the high level of synchronization prevents their representation in the MF formulation (Montbrió et al., 2015).

O_*P*_ and O_*S*_ emerge as sustained oscillations in the network thanks to a mechanism similar to that reported for pyramidal-interneuron gamma (PING) rhythms (Whittington et al., 2011) despite the frequency of these oscillations are not restricted to the *γ* band. Excitatory neurons start to fire followed by the inhibitory ones and the peak of activity of the excitatory population precedes that of the inhibitory one of a time delay Δ*t*. Furthermore, Δ*t* tends to vanish when the amplitude of the current fluctuations in the network increases. Indeed, for O_*F*_ oscillations, which cannot emerge in absence of current fluctuations, no delay has been observed between the activation of excitatory and inhibitory population.

A last important question that we tried to address in our work is if the COs are observable in the limit *N >> K >>* 1. We observed that the frequency of COs diverges as *K*^1*/*4^ with the median in-degree, while the amplitude of the COs appear to grow (vanish) with *K* for O_*F*_ (O_*P*_) oscillations in the examined range of in-degrees.

The paper is organized as follows. Section 2 is devoted to the introduction of the network model and of the corresponding effective neural mass model, as well as of the microscopic and macroscopic indicators employed to characterize the neural dynamics. In the same Section, the stationary solutions for the balanced neural mass model are analytically obtained as finite in-degree expansion and their range of stability determined together with bifurcation phase diagrams displaying the possible dynamical states. Network dynamics is studied in Section 3, in particular the asynchronous balanced state is analysed in details for structurally heterogeneous networks. Furthermore, the different types of COs observable in the balanced network with finite in-degrees are also characterized in Section 3. A brief summary of the results and conclusions is reported in Section 4.

## 2 MODELS AND DYNAMICAL INDICATORS

### 2.1 Network Model

We consider two sparsely coupled excitatory and inhibitory populations composed of *N* ^(*e*)^ and *N*^(*i*)^quadratic integrate-and-fire (QIF) neurons, respectively. (Ermentrout and Kopell, 1986). The evolution equation for the membrane potentials 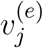 and 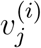 of the excitatory and inhibitory neurons can be written as:

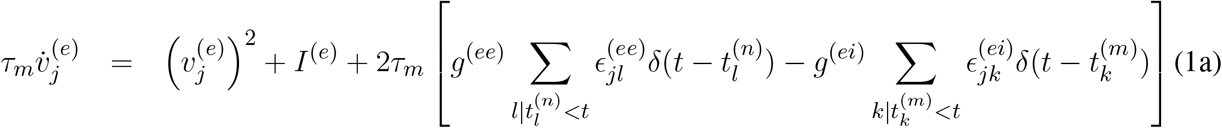

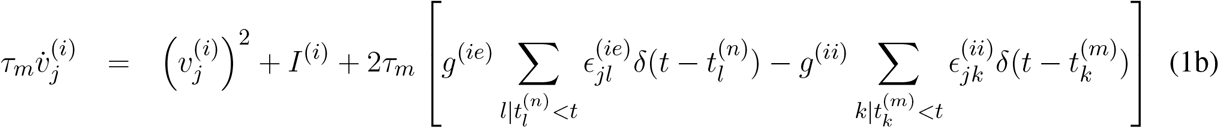

where *τ*_*m*_ = 20 ms is the membrane time constant that we set identical for excitatory and inhibitory neurons, *I*^(*e*)^ (*I*^(*i*)^) is the external DC current acting on excitatory (inhibitory) population, *g*^(*αβ*)^ represents the synaptic coupling strengths between post-synaptic neurons in population *α* and pre-synaptic ones in population *β*, with *α, β ∈{e, i}*. The elements of the adjacency matrices 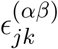 are equal to 1 (0) if a connection from a pre-synaptic neuron *k* of population *β* towards a post-synaptic neuron *j* of population *α*, exists (or not). Furthermore, 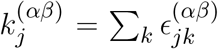 is the number of pre-synaptic neurons in population *β* connected to neuron *j* in population *α*, or in other terms its in-degree restricted to population *β*. The emission of the *n*-th spike emitted by neuron *l* of population *α* occurs at time 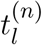 whenever the membrane potential 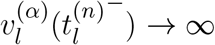, while the reset mechanism is modeled by setting 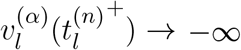 immediately after the spike emission. The post-synaptic potentials are assumed to be *δ*-pulses and the synaptic transmissions to be instantaneous. The equations (1) can be formally rewritten as

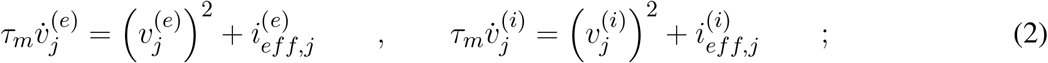

where 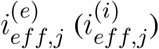 represents the instantaneous excitatory (inhibitory) effective currents, which includes the external DC current as well as the synaptic currents due to the recurrent connections.

We consider the neurons within excitatory and inhibitory population as randomly connected, with in-degrees *k*^(*αα*)^ distributed according to a Lorentzian distribution

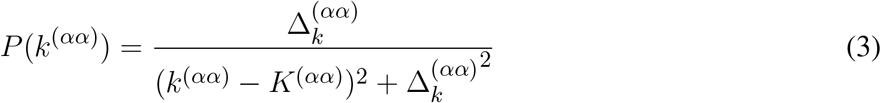

peaked at *K*^(*αα*)^ and with a half-width half-maximum (HWHM) 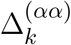, this latter parameter measures the level of structural heterogeneity in each population. For simplicity, we set *K*^(*ee*)^ = *K*^(*ii*)^ *≡K*. Furthermore, we assume that also neurons from a population *α* are randomly connected to neurons of a different population *β* ≠ *α*. However, in this case we consider no structural heterogeneity with in-degrees fixed to a constant value *K*^(*ei*)^ = *K*^(*ie*)^ = *K*. We have verified that by considering Erdös-Renyi distributed in-degrees *K*^(*ei*)^ and *K*^(*ie*)^ with average *K* does not modify the observed dynamical behaviour.

The DC current and the synaptic coupling are rescaled with the median in degree as 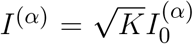 and 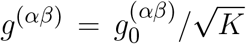, as done in previous works to obtain a self-sustained balanced dynamics for *N >> K >>* 1 (van Vreeswijk and Sompolinsky, 1996; Renart et al., 2010; Litwin-Kumar and Doiron, 2012; Kadmon and Sompolinsky, 2015). The structural heterogeneity parameters are rescaled as 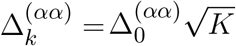 in analogy to Erdös-Renyi networks. The choice of the Lorentzian distribution for the *k*^(*αα*)^ is needed in order to obtain an effective MF description for the microscopic dynamics (di Volo and Torcini, 2018; Bi et al., 2020) as detailed in the next section.

The microscopic activity can be analyzed by considering the inter-spike interval (ISI) distribution as characterized by the coefficient of variation *cv*_*i*_ for each neuron *i*, which is the ratio between the standard deviation and the mean of the ISIs associated to the train of spikes emitted by the considered neuron. To characterize the macroscopic dynamics of each population *α* we measure the average coefficient of variation 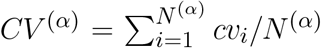, the mean membrane potential 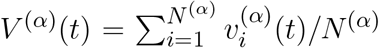 and the population firing rate *R*^(*α*)^(*t*), corresponding to the number of spikes emitted within population *α* per unit of time and per neuron.

Furthermore, the level of coherence in the neural activity of population *α* can be quantified in terms of the following indicator (Golomb, 2007)

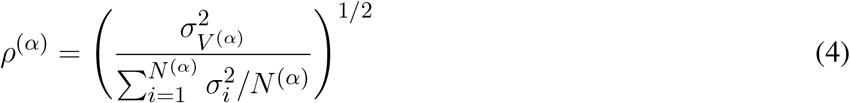

where 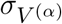 is the standard deviation of the mean membrane potential, 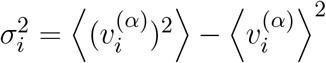 and ⟨*·* ⟩ denotes a time average. A perfect synchrony corresponds to *ρ*^(*α*)^ = 1, while an asynchronous dynamics to a vanishing small 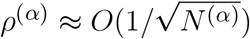.

The frequencies associated to collective motions can be identified by measuring the power spectra *S*(*ν*) of the mean membrane potentials *V* (*t*) of the whole network. In case of a periodic motion the position of the main peak *ν*_*CO*_ represents the frequency of the COs, while for quasi-periodic motions the spectrum is characterized by many peaks that can be obtained as a linear combination of two fundamental frequencies (*ν*_1_, *ν*_2_). The spectra obtained in the present case always exhibits also a continuous background due to the intrinsic fluctuations present in the balanced network. The power spectra have been obtained by calculating the temporal Fourier transform of *V* (*t*) sampled at time intervals of 10 ms. Time traces composed of 10000 consecutive intervals have been considered to estimate the spectra, which are obtained at a frequency resolution of Δ*ν* = 0.01 Hz. Finally, the power spectra have been averaged over five independent realizations of the random network.

The network dynamics is integrated by employing an Euler scheme with time step *dt* = 0.0001 ms, while time averages and fluctuations are usually estimated on time intervals *T*_*s*_ *≃* 100 s, after discarding transients *T*_*t*_ *≃* 10 s. Usually we consider networks composed of *N* ^(*e*)^ = 10000 excitatory and *N* ^(*i*)^ = 2500 inhibitory neurons.

### 2.2 Effective neural mass model

By following (Montbrió et al., 2015; di Volo and Torcini, 2018), we derive an effective neural mass formulation for the spiking network (1). As a starting point we consider an exact macroscopic model recently derived for a fully coupled homogeneous population of pulse-coupled QIF (Montbrió et al., 2015) with synaptic couplings randomly distributed according to a Lorentzian. The neural mass model corresponding to this QIF network can be written in terms of only two collective variables (namely, the mean membrane potential *V* and the instantaneous population rate *R*), as follows (Montbrió et al., 2015):

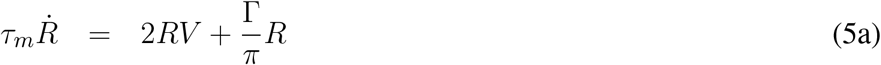

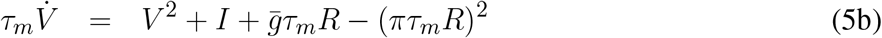

where 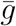 is the median and Γ the HWHM of the Lorentzian distribution of the synaptic couplings.

Such formulation can be applied to the sparse networks studied in this paper, indeed as shown in (di Volo and Torcini, 2018; Bi et al., 2020) for a single sparse inhibitory population the quenched disorder associated to the in-degree distribution can be rephrased in terms of random synaptic couplings. Namely, each neuron *i* in population *α* is subject to currents of amplitude 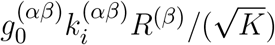 proportional to their in-degrees 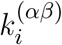, with *β* ∈ {*e, i*}. Therefore we can consider the neurons as fully coupled, but with random values of the couplings distributed as Lorentzian of median 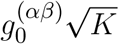 and HWHM 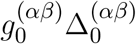. Since we have assumed a constant value

The neural mass model corresponding to the spiking network (1) can be written as follows:

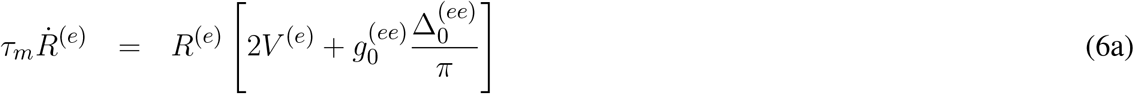

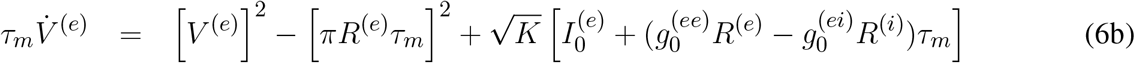

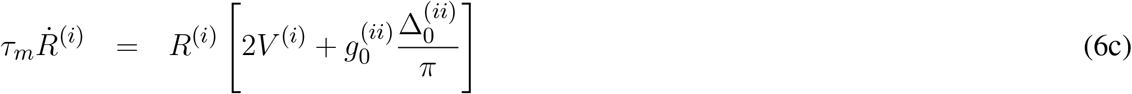

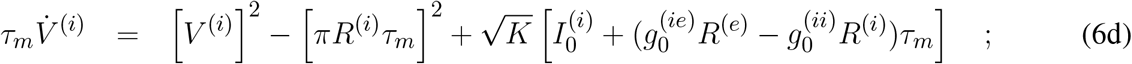

where we have set 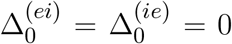, since we have assumed that the connections among neurons of different populations are random but with a fixed in-degree *K*^(*ei*)^ = *K*^(*ie*)^ = *K*.

#### 2.2.1 Stationary Solutions

The stationary solutions 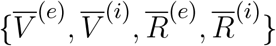 of (6) can be explicitly obtained for the mean membrane potentials as

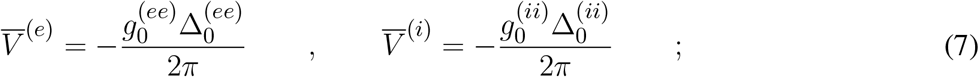

while the instantaneous population rates are the solutions of the following quadratic system

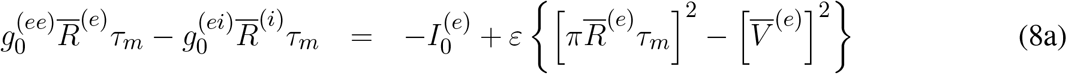

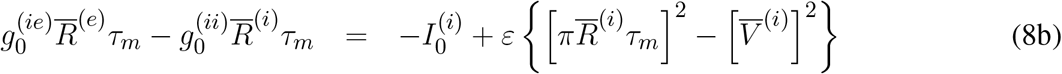

where 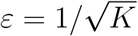 is a smallness parameter taking in account finite in-degree corrections. It is interesting to notice that the parameters controlling the structural heterogeneity 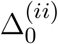 and 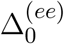 fix the stationary values of the mean membrane potentials reported in (7). The solutions of (8) can be exactly obtained and the associated bifurcations analysed by employing the software XPP AUTO developed for orbit continuation (Ermentrout, 2007).

For sufficiently large *K* one can obtain analytic approximations of the solution of (8) by expanding the population rates as follows

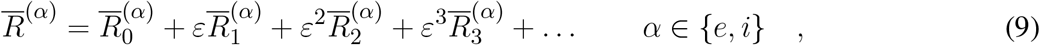

by inserting these expressions in (8), and finally by solving order by order in *ε*.

The solutions at any order can be written as follows:

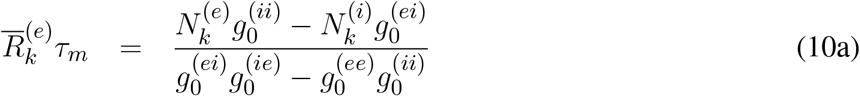

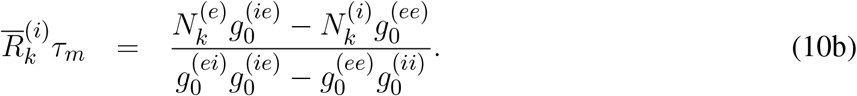

where

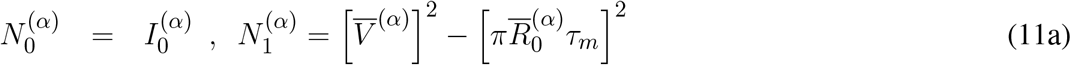

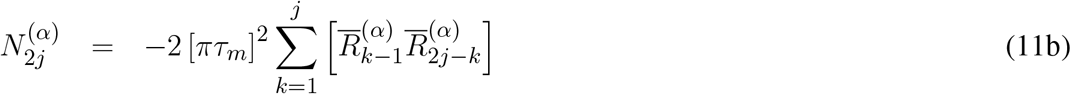

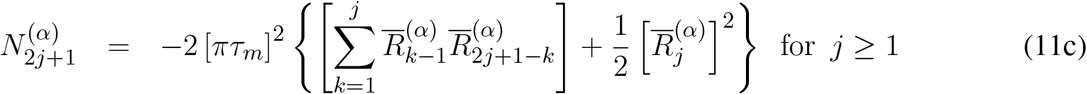

The systems (10) with parameters given by (11) can be resolved recursively for any order and the final solution obtained from the expression (9). The zeroth order approximation, valid in the limit *K → ∞*, corresponds to the usual solution found for rate models in the balanced state (van Vreeswijk and Sompolinsky, 1996; Rosenbaum and Doiron, 2014), such solution is physical whenever one of the following inequalities is satisfied

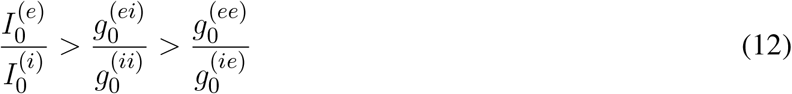

or

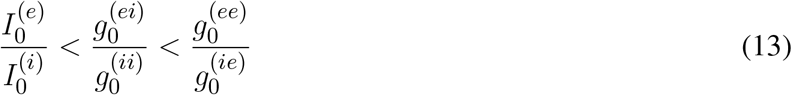

which ensure the positive sign of 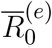 and 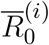. The zeroth order solution does not depend on the structural heterogeneity, since the ratio Δ^(*αα*)^*/K* vanishes in the limit *K → ∞*. It should be stressed that this ratio does not correspond to the coefficient of variation introduced in (Landau et al., 2016) to characterize the in-degree distribution. This because we are considering a Lorentzian distribution, where the average and the standard deviation are not even defined. Moreover, already the first order corrections depends on 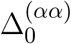.

In order characterize the level of balance in the system one usually estimates the values of the effective input currents 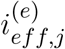 and 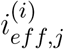 driving the neuron dynamics. These at a population level can be rewritten as

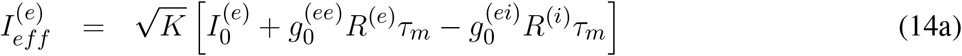

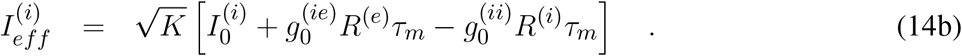

In a balanced state these quantities should not diverge with *K*, instead they should approach some constant value. In or MF formulation, we can estimate analytically the values of the effective currents in the limit *K → ∞* for an asynchronous state and they read as

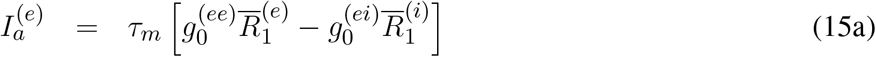

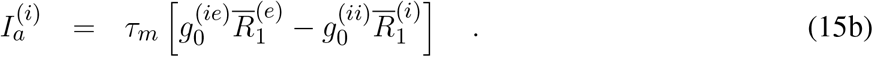

It should be noticed that these asymptotic values depend on the first order corrections to the balanced solution (10). Therefore, they depend not only on the synaptic couplings 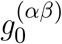 and on the external DC currents, but also on the parameters 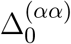 controlling the structural heterogeneities.

Depending on the parameter values, the currents 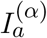 can be positive or negative, thus indicating a balanced dynamics where most part of the neurons are supra or below threshold, respectively. Usually, in order to obtain a stationary state characterized by a low rate and a Poissonian statistic, as observed in the cortex, one assumes that the excitation and inhibition nearly cancel. So that the mean membrane potential remains slightly below threshold, and the neurons can fire occasionally due to the input current fluctuations (van Vreeswijk and Sompolinsky, 1996; Brunel, 2000). However, as pointed out in (Lerchner et al., 2006) this is not the only possible scenario for a balanced state. In particular, the authors have developed a self-consistent MF theory for balanced Erdös-Renyi networks made of heterogeneous Leaky Integrate-and-Fire (LIF) neurons. In this context they have shown that Poisson-like dynamics are visible only at intermediate synaptic couplings. While mean driven dynamics are expected for low couplings, and at large couplings bursting behaviours appear in the balanced network. Recently, analogous dynamical behaviours have been reported also for a purely inhibitory heterogeneous LIF network (Angulo-Garcia et al., 2017). These findings are consistent with the results in (Lerchner et al., 2006), where the inhibition is indeed predominant in the balanced regime.

#### 2.2.2 Lyapunov analysis

To analyse the linear stability of generic solutions of Eqs. (6), we have estimated the corresponding Lyapunov spectrum (LS) *{λ*_*k*_*}* (Pikovsky and Politi, 2016). This can be done by considering the time evolution of the tangent vector *δ* = *{δR*(*e*), *δV* (*e*), *δR*(*i*), *δV* ^(*i*)^}, that is ruled by the linearization of the Eqs.(6), namely

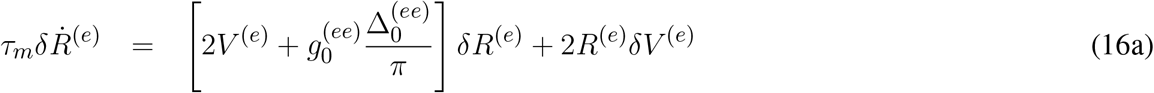

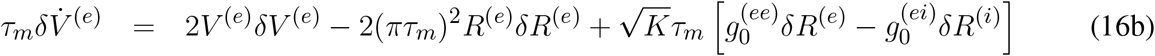

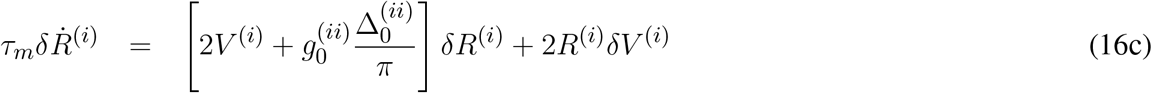

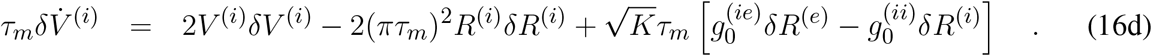

In this case, the LS is composed by four Lyapunov exponents (LEs) {*λ*_*k*_ } with *k* = 1, …, 4, which quantify the average growth rates of infinitesimal perturbations along the orthogonal manifolds. The LEs can be estimated as follows

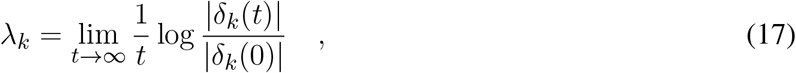

where the tangent vectors *δ*_*k*_ are maintained ortho-normal during the time evolution by employing a standard technique introduced in (Benettin et al., 1980). The autonomous system will be chaotic for *λ*_1_ *>* 0, while a periodic (two frequency quasi-periodic) dynamics will be characterized by *λ*_1_ = 0 (*λ*_1_ = *λ*_2_ = 0) and a fixed point by *λ*_1_ *<* 0.

In order to estimate the LS for the neural mass model we have integrated the direct and tangent space evolution with a Runge-Kutta 4th order integration scheme with *dt* = 0.01 ms, for a duration of 200 s, after discarding a transient of 10s

#### 2.2.3 Linear Stability of Stationary Solutions

The linear stability of the stationary solutions 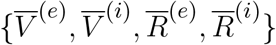 can be analyzed by solving the eigenvalue problem for the linear equations (16) estimated for stationary values of the mean membrane potentials and of the population firing rates. This approach gives rise to a fourth order characteristic polynomial of the complex eigenvalues 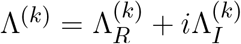 with *k* = 1, …, 4. The stability of the fixed point is controlled by the maximal 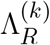, whenever it is positive (negative) the stationary solution is unstable (stable). The nature of the fixed point is determined by 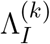, if the imaginary parts of the eigenvalues are all zero we have a node, otherwise a focus. Due to the fact that the coefficients of the characteristic polynomial are real the eigenvalues are real or if complex they appear in complex conjugates couples 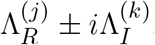. Therefore the relaxation towards the fixed point is characterized by one or two frequencies 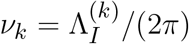. These latter quantities, as discussed in details in the following, can give good predictions for the frequencies *ν*_*CO*_ of fluctuation driven COs observable for the same parameters in the network dynamics.

In the limit *K >>* 1, we can approximate the linear stability equations (16) as follows:

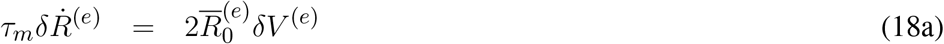

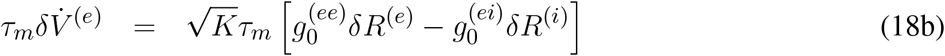

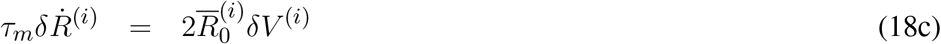

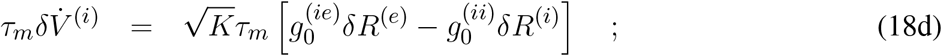

where we have considered the zeroth order approximation for the population rates 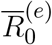 and 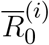.

In this case the complex eigenvalues Λ^(*k*)^ are given by the following expression:

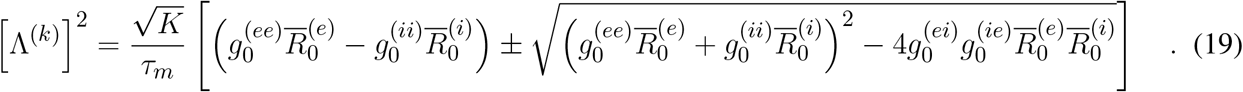

From (19) it is evident that Λ^(*k*)^ *∝* (*K*)^1*/*4^ and by assuming 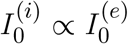, as we will do in this paper, we also have that 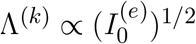. Therefore for a focus solution we will have the following scaling relation for the relaxation frequencies for sufficiently large *K*

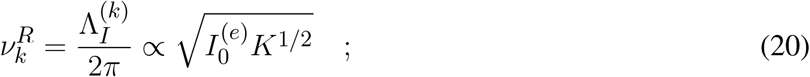

this scaling is analogous to that found for purely inhibitory QIF networks in (di Volo and Torcini, 2018). In (van Vreeswijk and Sompolinsky, 1996) it has been found that the eigenvalues, characterizing the stability of the asynchronous state, scale proportionally to 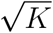, therefore the convergence (divergence) from the stationary stable (unstable) solution is somehow slower with *K* in our model. This is due to the presence in our MF of an extra macroscopic variable, the mean membrane potential, with respect to the usual rate models.

#### 2.2.4 Bifurcation diagrams

The dynamics of the neural mass model (6) takes place in a four dimensional space *{R*^(*e*)^, *V*^(*e*)^, *R*^(*i*)^, *V*^(*i*)^} and it depends on 9 parameters, namely on the four synaptic coupling strengths 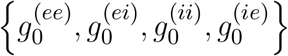, the two external stimulation currents 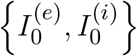, the median in-degree *K* and the HWHM of the two distributions of the in-degrees 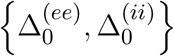.

However, in order to reduce the space of parameters to investigate and at the same time to satisfy the inequalities (12), required for the existence of a balanced state in the large *K* limit, we fix the inhibitory DC current as 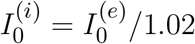 and the synaptic couplings as 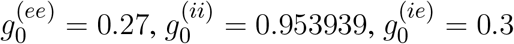, and 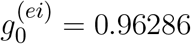 analogously to what done in (Monteforte and Wolf, 2010). Therefore we are left with four control parameters, namely 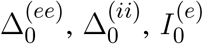, and *K*, that we will vary to investigate the possible dynamical states.

Three bidimensional bifurcation diagrams for the neural mass model (6) are reported in Fig. 1 for the couples of parameters 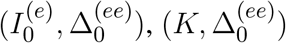 and 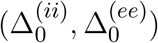. From the bifurcation analysis we have identified five different dynamical states for the excitatory population : namely, (I) an unstable focus; (II) a stable focus coexisting with an unstable limit cycle; (III) a stable node; (IV) a stable limit cycle coexisting with an unstable focus; (V) a chaotic regime. For the analysis reported in the following it is important to remark that the stable foci are usually associated to four complex eigenvalues arranged in complex conjugate couples, therefore the relaxation towards a stable focus is characterized by two frequencies (*ν*_1_, *ν*_2_) corresponding to the complex parts of the eigenvalues. In region (III) the macroscopic fixed point is characterized by two real eigenvalues and a couple of complex conjugated ones. Thus the relaxation towards the macroscopic node is in this case guided by a single relaxation frequency. The inhibitory population reveals the same bifurcation structure as the excitatory one, apart an important difference: the inhibitory population never displays stable nodes. Therefore the region (III) for the inhibitory population is also a region of type (II).

**Figure 1.**
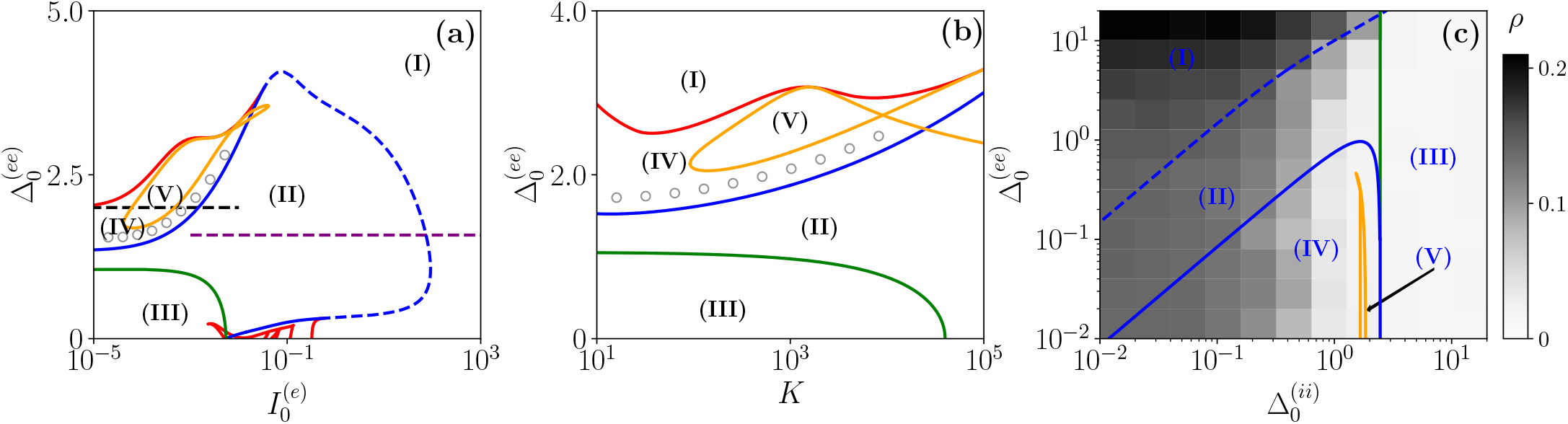
Bifurcation diagrams of the neural mass model. The bifurcation diagrams concerns the dynamical state exhibited by the excitatory population in the bidimensional parameter spaces 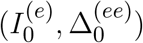 (a), 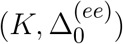 (b) and 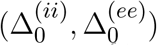. The regions marked by Roman numbers correspond to the following collective solutions: (I) an unstable focus; (II) a stable focus coexisting with an unstable limit cycle; (III) a stable node; (IV) an unstable focus coexisting with a stable limit cycle; (V) a chaotic dynamics. The green solid line separates the regions with a stable node (III) and a stable focus (II). The blue solid (dashed) curve is a line of super-critical (sub-critical) Hopf bifurcations (HBs), and the red one of Saddle-Node (SN) bifurcations of limit cycles. The yellow curve denotes the period doubling (PD) bifurcation lines. In (c) we report also the coherence indicator *ρ*^(*e*)^ (4) estimated from the network dynamics with *N* ^(*e*)^ = 10000 and *N* ^(*i*)^ = 2500. The dashed lines in (a) indicate the parameter cuts we will consider in Figs. 4 and 5 (black) and Fig. 7 (purple), while the open circles in (a) and (b) denote the set of parameters employed in Fig. 11. In the three panels the inhibitory DC current and the synaptic couplings are fixed to 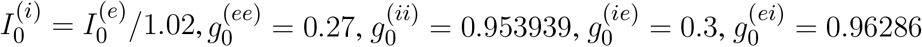; other parameters: (a) *K* = 1000, 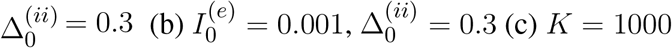 and 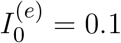.

As shown in Fig. 1 (a) and (b), for fixed 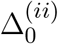 and for low values of the structural heterogeneity 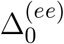 and of the excitatory DC current 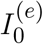 one observes a stable node (III) that becomes a stable focus (II) by increasing 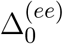, these transitions are signaled as green solid lines in Fig. 1. By further increasing the degree of heterogeneity 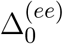, the stable focus gives rise to collective oscillations (IV) via a super-critical Hopf Bifurcation (HB) (blue solid lines). Depending on the values of *K* and 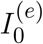 one can have the emergence of chaotic behaviours (V) via a period doubling (PD) cascade (yellow solid lines). For sufficiently large 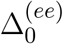, the COs disappear via a Saddle-Node (SN) bifurcation of limit cycles (red solid lines) and above the SN line the only remaining solution is an unstable focus (I).

As shown in Fig. 1 (a), for fixed structural heterogeneities the increase of 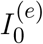 leads to the disappearance of the stable focus (II) via a sub-critical HB (dashed blue line). The dependence of the observed MF solutions on the in-degree *K* is reported in Fig. 1 (b) for a current 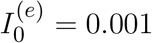 and it is not particularly dramatic, apart for the emergence of a chaotic region (V) from a CO regime (IV).

In order to observe the emergence of COs (IV) from the destabilization of a node solution (III) we should vary the structural inhibitory heterogeneity 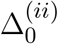, as shown in Fig. 1 (c). Indeed, for sufficiently low 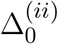 and 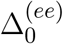 we can observe super-critical bifurcation line from a node to a stable limit cycle (LC). From this analysis it emerges that the excitatory heterogeneity has an opposite effect with respect to the inhibitory one, indeed by increasing 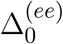 the value of *ρ*^(*e*)^ increases indicating the presence of more synchronized COs. This effect is due to the fact that the increase of 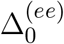 leads to more and more neurons with large 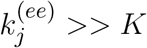, therefore receiving higher and higher levels of recurrent excitation. These neurons are definitely supra-threshold and drive the activity of the network towards coherent behaviours.

In order to understand the limits of our MF formulation, it is of particular interest to compare the network simulations with the MF phase diagram. To this aim, we report in Fig. 1 (c) also the the coherence indicator *ρ*^(*e*)^ (4) estimated from the network dynamics. The indicator *ρ*^(*e*)^ reveals that no COs are present in region (III), where the MF displays a stable node, however COs emerge in all the other MF regimes for sufficiently low 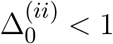. The presence of COs is expected from the MF analysis only in the regions (IV) and (V), but neither in (II) where the MF forecasts the existence of a stable focus nor in (I) where no stable solutions are envisaged. The origin of the discrepancies among the MF and the network simulations in region (II) is due to the fact that the considered neural mass neglects the dynamical fluctuations in the input currents present in the original networks, that can give rise to noise induced COs (Goldobin et al., 2021). However, as shown in (di Volo and Torcini, 2018; Bi et al., 2020) for purely inhibitory populations, the analysis of the neural mass model can still give relevant information on the network dynamics. In particular, the frequencies of the fluctuation induced COs observable in the network simulations can be well estimated from the frequencies (*ν*_1_, *ν*_2_) of the relaxation oscillations towards the stable MF focus. The lack of agreement between MF and network simulations in the region (I) is due to finite size effects, indeed in this case the system tends to fully synchronize. Therefore, in the network one observes highly synchronized COs characterized by population firing rates that diverge for increasing *K* and *N* and the MF is unable to reproduce these unrealistic solutions (Montbrió et al., 2015).

On the basis of these observations, we can classify the COs observable in the network in three different types accordingly to the corresponding MF solutions: O_P_, when in the MF we observe periodic, quasi-periodic or chaotic collective solutions in regions (IV) and (V); O_F_, when the MF displays relaxation oscillations towards the stable focus in regions (II) and (III), that in the sparse network become noise sustained oscillations due to fluctuations in the input currents; O_S_, when the MF fully synchronizes as in region (I).

In the following we will investigate in details the network dynamics for asynchronous regimes, as well as for two main scenarios indicated as dashed horizontal lines in Fig. 1 (a), where COs are observable, corresponding to the transition to chaos (black dashed line) and to the emergence of abnormal synchronization from a stable focus (purple dashed line).

## 3 NETWORK DYNAMICS

In this section we analyse the dynamics of the E-I network of QIF neurons with particular emphasis on the macroscopic behaviours in order to test for the predictions of the effective neural mass model. In particular, we will consider asynchronous as well as coherent dynamics, in this latter case we will focus on the three types of identified COs: namely, O_P_, O_F_ and O_S_. These can manifest as periodic, quasi-periodic and chaotic solutions as we will see in the following.

### 3.1 Asynchronous Regimes

We will firstly consider a situation where the network dynamics remains asynchronous for any value of the median in-degree *K*, this occurs for sufficiently high structural inhibitory heterogeneities 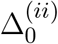 and external DC currents as shown in Fig. 1 (b) and (c) for E-I networks and as reported in (di Volo and Torcini, 2018) for purely inhibitory populations. If the population dynamics is asynchronous, we expect that at a MF level the system will converge towards a stationary state corresponding to a stable equilibrium. Therefore we have compared the results of the network simulations with the stationary rates 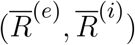 solutions of (6). As shown in Fig. 2 (a) and (b), the macroscopic activity of the excitatory and inhibitory populations is well reproduced by the fixed point solutions (8) in a wide range of values of the in-degrees 10 *≤ K ≤* 10^4^. This is particularly true for the inhibitory population, while at low *K <* 100 the excitatory firing rate is slightly underestimate by the macroscopic solution 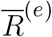. Due to our choice of parameters, the average inhibitory firing rate is definitely larger than the excitatory one for *K >* 100. This is consistent with experimental data reported for the barrel cortex of behaving mice (Gentet et al., 2010) and other cortical areas (Mongillo et al., 2018). Moreover, the rates have a non monotonic behaviour with *K* with a maximum at *K ≃* 450 (*K ≃* 2500) for excitatory (inhibitory) neurons. As expected, the balanced state solutions 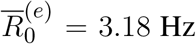 and 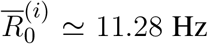 (dashed horizontal lines) are approached only for sufficiently large *K >>* 1. In Fig. 2 (a) and (b) are reported also the first (second) order approximation 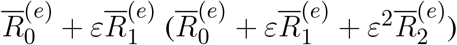 given by Eq. (10). These approximations reproduce quite well the complete solutions already at *K ≥* 10^4^.

**Figure 2.**
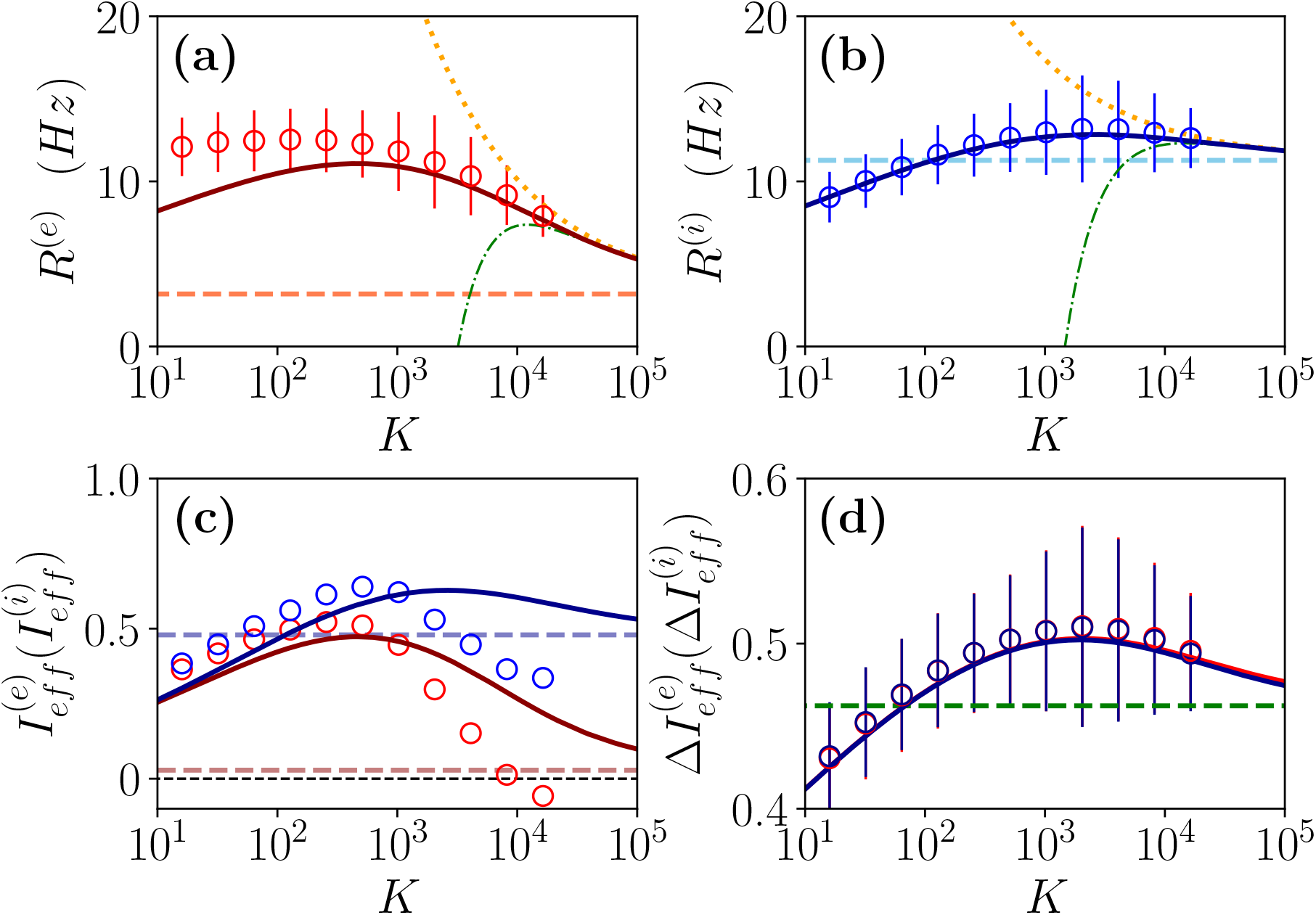
Asynchronous Dynamics. Instantaneous population rate *R*^(*e*)^ (*R*^(*i*)^) of excitatory (inhibitory) neurons in function of the median in-degree *K* are shown in panel (a) (panel (b)). The effective input currents 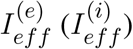 given by Eqs. (14) are reported in panel (c) and the fluctuations of the input currents 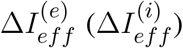, as obtained from Eqs. (21), in panel (d). Red (blu) color refer to excitatory) inhibitory population. The solid continuous lines represent the value obtained by employing the exact MF solutions 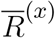 of (8), the dotted (dash-dotted) lines correspond to the first (second) order approximation 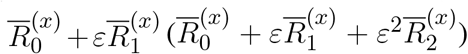 and the dashed horizontal lines to the zeroth order one 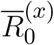 in (a),(b) and (d), and to 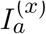 in (c) with *x* = *e, i*. The circles correspond to data obtained from numerical simulations of *N* ^(*e*)^ = *N* ^(*i*)^ = 10000 neurons for *K <* 4096, *N* ^(*e*)^ = *N* ^(*i*)^ = 20000 for *K* = 4096, 8192 and *N* ^(*e*)^ = *N* ^(*i*)^ = 30000 for *K >* 8192, averaging the population rates over a window of *T* = 40*s*, after discarding a transient of *T* = 60*s*. The error bars in (a) and (b) are obtained as the standard deviations (over the time window *T*) of the population rates, while the average CV of neurons is around 0.15 for all the reported simulations. Synaptic couplings and the ratio between the currents are fixed as stated in sub-section 2.2.4, other parameters are 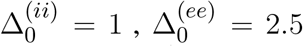 and 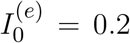. The values of the asymptotic solutions (dashed lines) are : in (a) and (b) 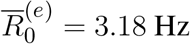 and 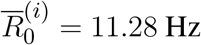, respectively; in (c) 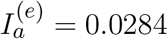 and 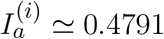; in (d) 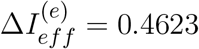 and 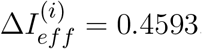.

Let us now consider the effective input currents (14), these are reported in Fig. 2 (c) versus the median in-degree. As expected, for increasing *K* the MF estimations of the effective currents (solid lines) converge to the asymptotic values 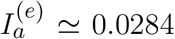 and 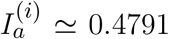 (dashed lines) for our choice of parameters. For the excitatory population the asymptotic value of the effective input current is essentially zero, while for the inhibitory population it is definitely positive. These results suggest that for the considered choice of parameters the dynamics of both populations will be balanced, since the quantities 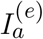 and 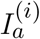 do not diverge with *K*, however at a macroscopic level the excitatory population will be at threshold, while the inhibitory one will be supra-threshold. For comparison, we have estimated 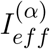 also from the direct the network simulations (circles) for 16 *≤K ≤*16384. These estimations disagree with the MF results already for *K >* 1000. This despite the fact that the population firing rates in the network are very well captured by the MF estimations at large *K*, as shown in Fig. 2 (a) and (b). These large differences in the effective input currents are clearly the effect of small discrepancies at the level of firing rates enhanced by the multiplicative factor 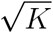 appearing in Eqs. (14). However, from the network simulations we observe that the effective currents approach values smaller than the asymptotic ones 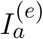 and 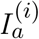 obtained from the the neural mass model. In particular, despite the fact that from finite *K* simulations it is difficult to extrapolate the asymptotic behaviours, it appears that 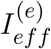 approaches a small negative value for *K >>* 1, while 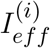 converges to some finite positive value. In the following we will see the effect of these different behaviours on the microscopic dynamics. The origin of the reported discrepancies should be related to the presence of current fluctuations in the network that are neglected in the MF formulation.

The relevance of the current fluctuations for the network dynamics can be appreciated by estimating their amplitudes within a Poissonian approximation, as follows

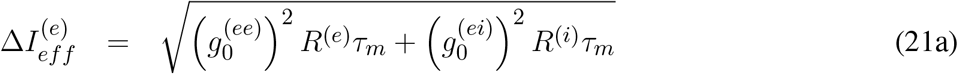

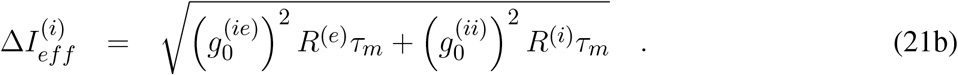

These have been evaluated by assuming that each neuron receive on average *K* excitatory and inhibitory spike trains characterized by a Poissonian statistics with average rates *R*^(*e*)^ and *R*^(*i*)^. However, we have neglected in the above estimation the variability of the in-degrees of each neuron. As shown in Fig. 2 (d), these fluctuations are essentially identical for excitatory and inhibitory neurons and coincide with the MF results. In the limit *K >>* 1 they converge to the asymptotic values 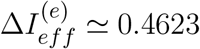 and 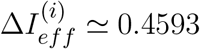 (green dashed lines). It is evident that already for *K >* 1000 the amplitudes of the fluctuations are of the same order or larger than the effective input currents. Thus suggesting that the fluctuations have indeed a relevant role in determining the network dynamics and that one would observes Poissonian or sub-Poissonian dynamics for the neurons, whenever 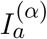 is sub-threshold or supra-threshold (Lerchner et al., 2006).

In order to understand how the in-degree heterogeneity influences the network dynamics at a microscopic level we examine the dynamics of active neurons in function of their total in-degree 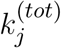. This is defined for excitatory (inhibitory) neurons as 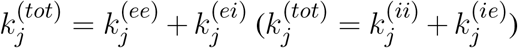. Furthermore, a neuron is considered as active if it has fired at least once during the whole simulation time *T*_*t*_ + *T*_*s*_ = 100 s, therefore if it has a firing rate larger than 0.01 Hz. As shown in Figs. 3 (a) and (b), the PDF of active neurons is skewed towards values 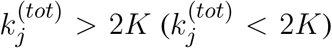 for excitatory (inhibitory) neurons. These results reflect the fact that the excitatory (inhibitory) neurons with low (high) recurrent in-degrees 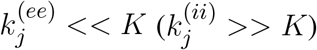 are driven below threshold by the inhibitory activity, that is predominant in the network since*R*^(*i*)^ *> R*^(*e*)^, 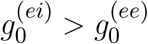, and 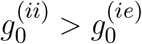. The number of silent neurons for *K >* 1024 is of the order of 6-10 % for both inhibitory and excitatory populations, in agreement with experimental results for the barrel cortex of mice (O’Connor et al., 2010), where a fraction of 10 % of neurons was identified as silent with a firing rate slower than 0.0083 Hz. It should be remarked that all the population averages we report include the silent neurons.

**Figure 3.**
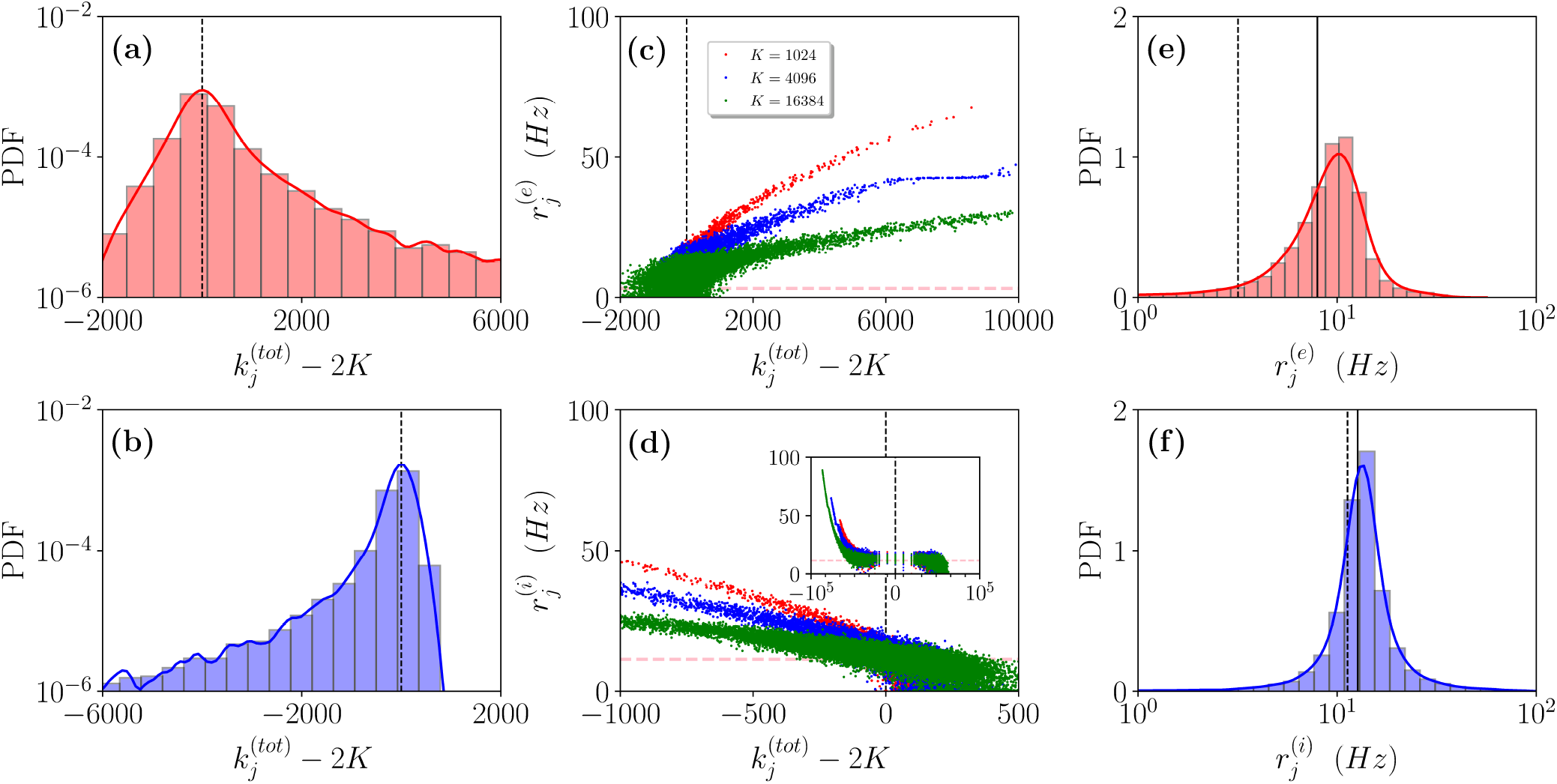
Asynchronous Dynamics. Probability distribution functions (PDFs) of the total in-degrees 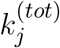 for excitatory (a) and inhibitory (b) active neurons for *K* = 16384. (c-d) Firing rates of the excitatory (inhibitory) neurons 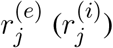 versus their total in-degrees 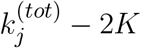 symbols refer to *K* = 1024 (red), *K* = 4096 (blue) and *K* = 16384 (green). The inset in (d) is an enlargement of the panel displaying the firing rates over the entire scale 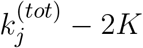. The magenta dashed lines in (c-d) represent the balanced state solution 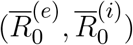. (e-f) PDF of the excitatory (inhibitory) firing rates 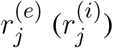 for *K* = 16384, the solid (dashed) line refers to the MF results 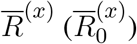 with *x* = *e, i*. The red (blue) solid line refers to a log-normal fit to the excitatory (inhibitory) PDF with mean 8.8 Hz (17.5 Hz) and standard deviation 3.8 Hz (2.3 Hz). The parameters are the same as in Fig. 2, the firing rates have been estimated by simulating the networks for a total time *T*_*s*_ = 60 s, after discarding a transient *T*_*t*_ = 40 s.

Let us now examine how the firing rates of active neurons will modify by increasing the value of the median in-degree *K*. The single neuron firing rates as a function of their total in-degrees 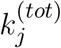 are reported in Figs. 3 (c) and (d) for *K* = 1024, 4096 and 16384. A common characteristics is that the bulk neurons, those with 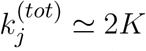, tend to approach the firing rate values 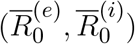 (magenta dashed lines) corresponding to the expected solutions for a balanced network in the limit *N >> K → ∞* (van Vreeswijk, 1996). This is confirmed by the analysis of their coefficient of variations *cv*_*j*_, whose values are of order one, as expected for fluctuation driven dynamics. On the other hand, the outlier neurons, i.e. those with 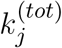 far from 2*K*, are all characterized by low values of the coefficient of variation *cv*_*j*_ indicating a mean driven dynamics. However, there is striking difference among excitatory and inhibitory neurons. For the excitatory ones we observe that the firing rates of the outliers with 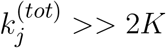 decrease for increasing *K*, while for the inhibitory population the increase of *K* leads to the emergence of outliers at 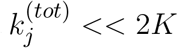 with higher and higher firing rates (see the inset in Fig.3 (d)). This difference can be explained by the different values measured for 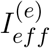 and 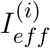 in the network (see Fig. 2 (c)). The increase of *K* leads for the excitatory (inhibitory) population to the emergence of neurons with very large 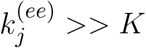 (very small 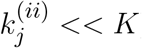) whose dynamics should be definitely supra-threshold. However, this is compensate in the excitatory case by the rapid drop of 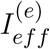 towards zero or negative values, while for the inhibitory population 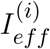 remains positive even at the largest *K* we have examined.

These outliers seem to have a negligible influence on the population dynamics, as suggested by the fact that the mean firing rates are well reproduced by the balanced solutions 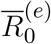 and 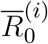 and as confirmed also by examining the PDFs of the firing rates for *K* = 16384. As shown in Figs. 3 (e) and (f), the excitatory (inhibitory) PDF can be well fitted by a log-normal distribution with mean 8.8 Hz (17.5 Hz) and standard deviation 3.8 Hz (2.3 Hz). This is considered a clear indication that the network dynamics is fluctuation driven (Roxin et al., 2011; Mongillo et al., 2018). However, the relative widths of our distributions are narrower than those reported in (Mongillo et al., 2018). This difference can find an explanation in the theoretical analysis reported in (Roxin et al., 2011), where the authors have shown that quite counter intuitively a wider distribution of the synaptic heterogeneities can lead to a narrower distribution of the firing rates. Indeed, here we consider Lorentzian distributed in-degrees, while in (Mongillo et al., 2018) Erdös-Renyi networks have been analyzed. As a further aspect, we have estimated the number of inhibitory neurons firing faster than a certain threshold *ν*_*th*_, this number does not depend on the median in-degree for sufficiently large *K >* 5000, however it grows proportionally to *N*. In the considered cases, the fraction of these neurons is *≃* 1% for *ν*_*th*_ = 50 Hz.

From this analysis we can expect in the limit *K >>* 1 to observe at a macroscopic scale an essentially balanced regime sustained by the bulk of active neurons, whose dynamics is fluctuation driven. However, at finite *K* and for finite observation times we will have a large body of silent neurons as well as a small fraction of mean driven outliers. These should be considered as typical features of finite heterogeneous neural circuits as shown in various experiments (O’Connor et al., 2010; Landau et al., 2016). Moreover, in the present case we report a quite different behaviours for outliers whose macroscopic effective input currents are supra or sub-threshold.

### 3.2 Collective Oscillations

We will now characterize the different type of COs observable by firstly following a route to coherent chaos for the E-I balanced network and successively we will examine how oscillations exhibiting an abnormal level of synchronization, somehow similar to those observable during an ictal state in the brain (Lehnertz et al., 2009), can emerge in our system. Furthermore, we will consider the phenomenon of quasi-periodicity and frequency locking occurring for fluctuation driven oscillations. As a last issue, the scaling of the frequencies and amplitudes of COs with the in-degree and as a function of the external DC current is reported.

#### 3.2.1 A period doubling route to coherent chaos

As a first case we will follow the path in the parameter space denoted as a dashed black line in Fig. 1 (a). In particular, in order to characterize the different dynamical regimes we have estimated the Lyapunov spectrum {*λ*_*i*_*}* associated to the MF equations. As shown in Fig. 4, this analysis has allowed to identify a period doubling cascade towards a chaotic region, characterized by periodic and chaotic windows. In particular, we observe a focus region (II) for 0.0015 < 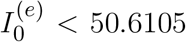, the focus looses stability via a super-critical Hopf bifurcation at 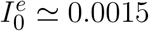 giving rise to COs. One observes a period doubling cascade (regime (V)) taking place in the interval 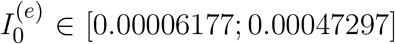 followed by a regime of COs at lower values of 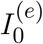. The chaotic dynamics refer to the MF evolution and it can be therefore definitely identified as collective chaos (Nakagawa and Kuramoto, 1993; Shibata and Kaneko, 1998; Olmi et al., 2011). A peculiar aspect of this period doubling cascade is that the chaotic dynamics remain always confined in four distinct regions without merging in an unique interval as it happens e.g. for the logistic map at the Ulam point (Ott, 2002). This is due to the fact that the population dynamics displays period four oscillations characterized by four successive bursts, whose amplitudes (measured by 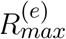) varies chaotically but each one remains restricted in an interval not overlapping with the other ones.

**Figure 4.**
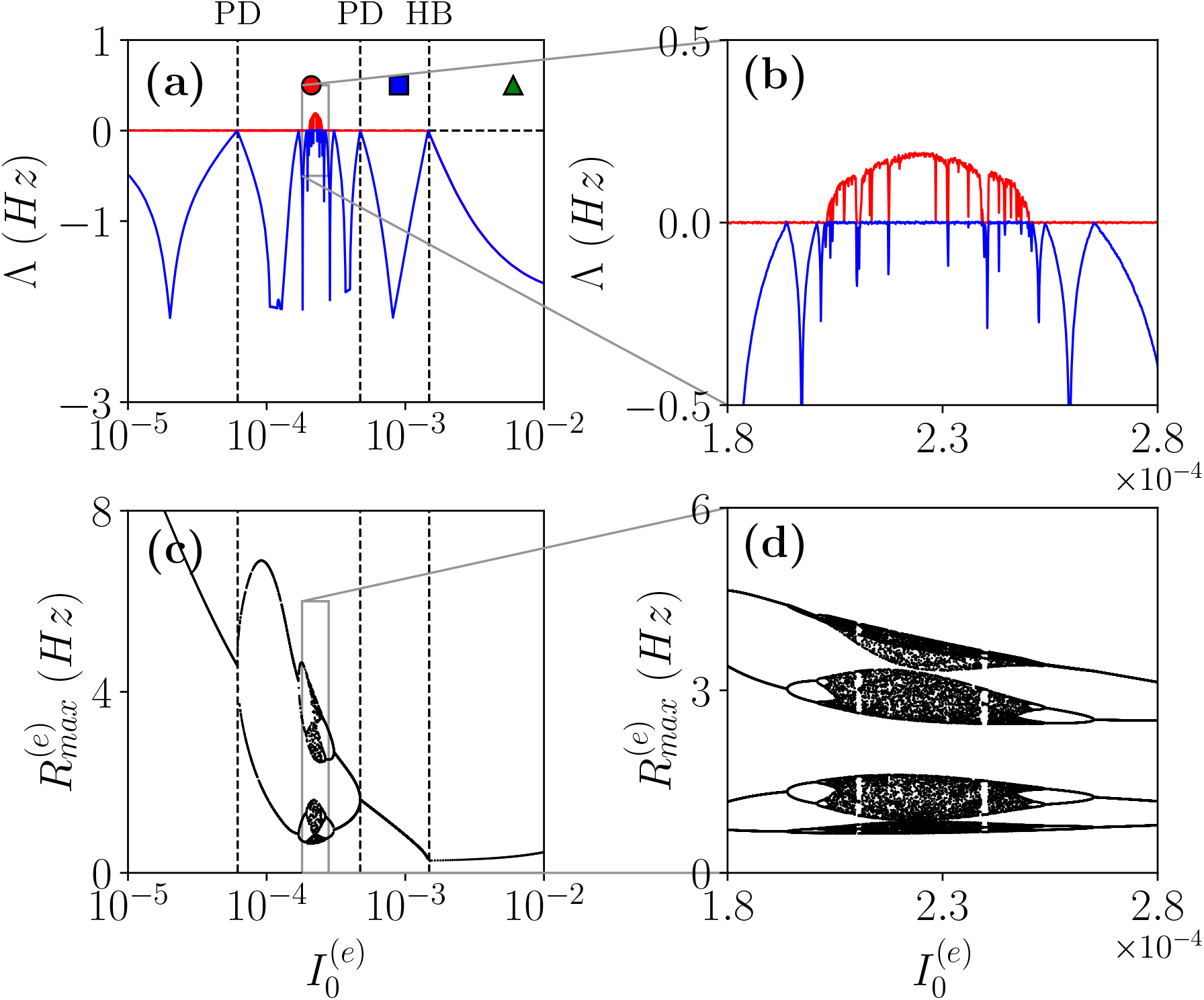
Coherent Chaos. (a-b) First (red) *λ*_1_ and second *λ*_2_ (blue) Lyapunov exponents for the MF versus the DC current 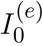 for the parameter cut corresponding to the dashed black line in Fig. 1 (a). The dashed vertical lines in (a) indicate a super-critical Hopf Bifurcation (HB) from a stable focus to periodic COs and the region of the Period Doubling (PD) cascade. The symbols denote three different types of MF solutions: namely, stable focus (green triangle); periodic oscillations (blue square) and chaotic oscillations (red circle). (c-d) Bifurcation diagrams for the same region obtained by reporting the maximal value of the instantaneous firing rate *R*^(*e*)^ measured from MF simulations. The parameters are the same as in Fig. 1, other parameters set as 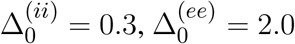, *K* = 1000.

Let us now examine the network dynamics for the 3 peculiar MF solutions indicated in Fig. 4 (a) corresponding to a stable focus (II) characterized by Lyapunov exponents (*λ*_1_ = *λ*_2_ = *−*0.0299, *λ*_3_ = *λ*_4_ = *−*0.101) for 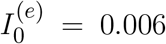 (green triangle), to a stable oscillation (IV) with (*λ*_1_ = 0.0, *λ*_2_ = *−*0.0343, *λ*_3_ = *−*0.0555, *λ*_4_ = *−*0.1732) for 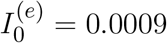 (blue square), and to collective chaos (v) with (*λ*_1_ = 0.0033, *λ*_2_ = 0.0, *λ*_3_ = *−* 0.0809, *λ*_4_ = *−* 0.1855) for 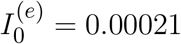 (red circle). As shown, in Fig. 5 in the network the dynamics is always characterized by oscillations for all the three considered cases: namely, O_P_ for the regimes (IV) and (V) and fluctuation induced O_F_ for to the stable MF focus.

**Figure 5.**
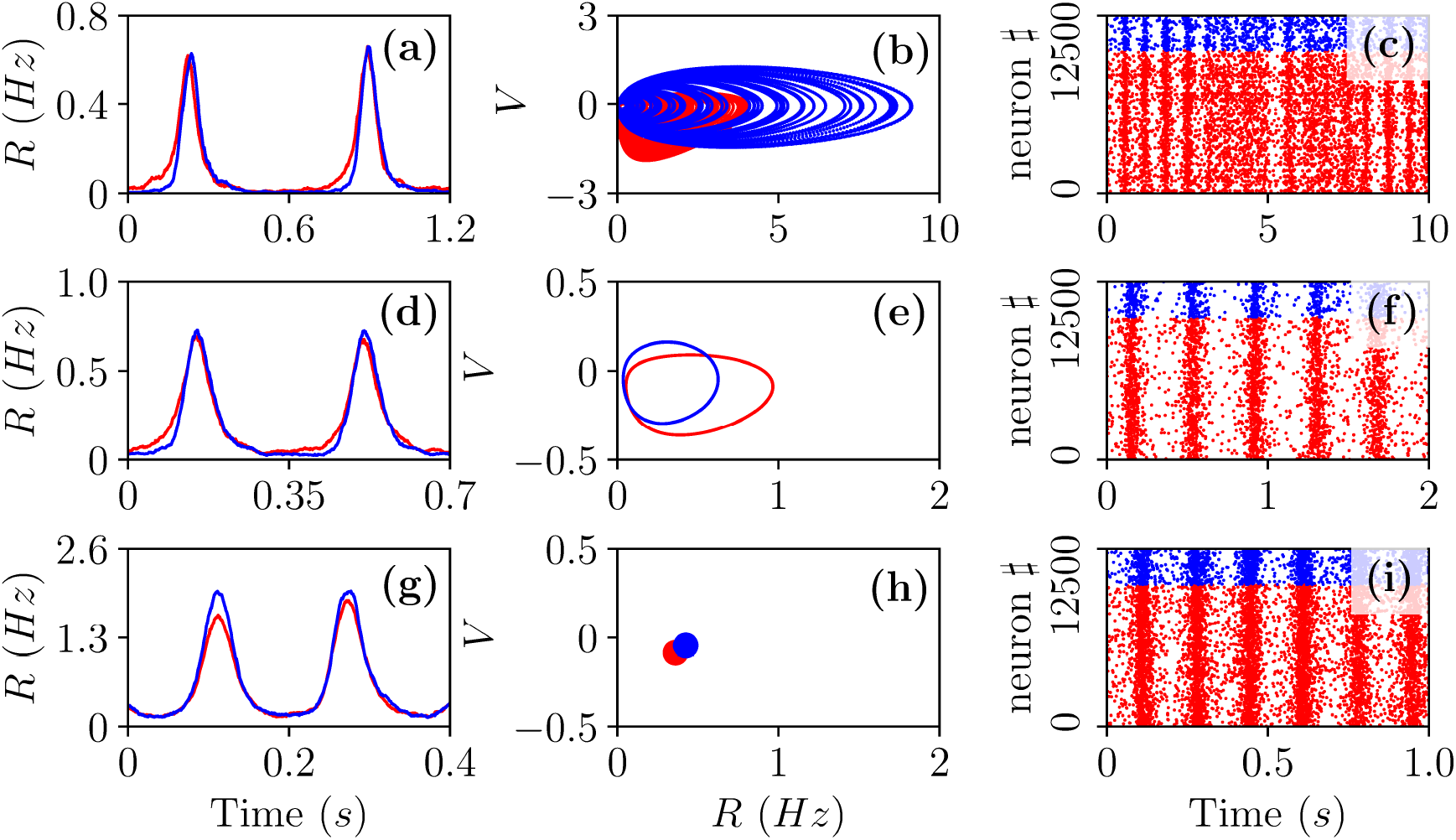
Different Types of Collective Oscillations. Panels (a-c) refer to the chaotic state observable for 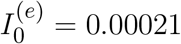 in the MF denoted by a red circle in Fig. 4 (a); panels (d-f) to the oscillatory state of the MF observable for 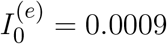 denoted by a blue square in Fig. 4 (a); panels (g-i) to the stable focus for the MF observable for 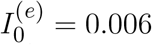 denoted by a green triangle in Fig. 4 (a). In the first (third) column are displayed the population firing rates (raster plots) versus time obtained from the network dynamics, while in the second column the corresponding MF attractors are reported in the planes identified by (*R*^(*e*)^, *V* ^(*e*)^) and (*R*^(*i*)^, *V* ^(*i*)^). Red (blue) color refers to excitatory (inhibitory) populations. Parameters as in Fig. 2, apart 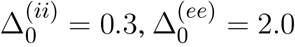, *K* = 1000. For the estimation of the firing rates we employed *N* ^(*e*)^ = 40000 and *N* ^(*i*)^ = 10000, while for the raster plots *N* ^(*e*)^ = 10000 and *N* ^(*i*)^ = 2500.

A typical feature of the O_P_ oscillations is that the excitatory neurons start to fire followed by the inhibitory ones, furthermore the peak of activity of the excitatory population usually precedes that of the inhibitory neurons of a time interval Δ*t*. Then the inhibitory burst silences the excitatory population for the time needed to recover towards the firing threshold. This recovering time sets the frequency *ν*_*CO*_ of the COs. In our set-up the excitatory bursts are wider than the inhibitory ones due to the fact that 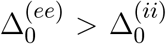.All these features are quite evident from the population firing rates shown in in Fig. 5 (a) and (d) and the raster plots in panel (c) and (f). These are typical characteristics of a PING-like mechanism reported for the generation of *γ* oscillations in the cortex (Tiesinga and Sejnowski, 2009), despite the fact that the CO’s frequencies shown in panels (a) and (d) are of the order of few Hz. Fluctuation driven oscillations O_F_ emerging in the network are radically different, as shown in Fig. 5 (g) in this case the excitatory and inhibitory populations deliver almost simultaneous bursts.

In order to understand the different mechanisms at the basis of O_P_ and O_F_ oscillations, let us examine how the delay Δ*t* between excitatory and inhibitory bursts, observed for O_P_ oscillations, modifies as a function of the membrane time constant of the inhibitory population 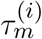. An increase of 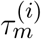 of *≃* 5 ms has the effect of reducing the delay of almost a factor six from Δ*t ≃* 28 ms to Δ*t ≃* 5 ms, as shown in Fig. 6 (a). The increase of 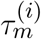 leads to an enhanced inhibitory action, since the integration of the inhibitory membrane potentials occurs on longer time scales and this promotes a higher activity of the inhibitory population. Indeed, this is confirmed from the drop of the effective input currents from an almost balanced situation where the average 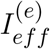 and 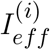 are almost zero to a situation where they are definitely negative (see Fig. 6 (b)). Thus for increasing 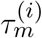 the percentage of neurons below threshold also increases and as a consequence the dynamics becomes more and more noise driven, as testified by the increase of the current fluctuations 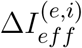 shown in Fig. 6 (c). In summary, the delay is due to the fact that, despite the effective inhibitory and excitatory currents are essentially equal, as shown in Fig. 6 (b), the wider distribution of the excitatory in-degrees promote the presence of excitatory neurons supra-threshold that are the ones igniting the excitatory burst before the inhibitory one. The delay Δ*t* decreases whenever the number of these supra-threshold neurons decreases and it will vanish when the dynamics will become essentially fluctuation driven as in the case of O_F_ oscillations.

**Figure 6.**
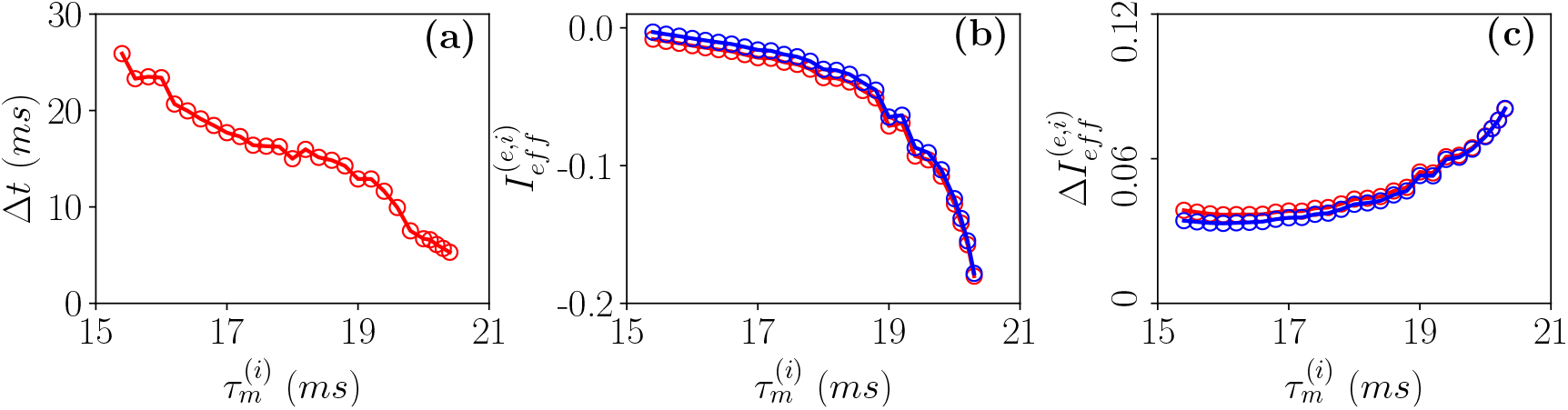
PING-like O_*P*_ collective oscillations. (a) Firing delays Δ*t* between the excitatory population peak and the inhibitory one versus 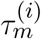. Effective mean input currents (14) (b) and current fluctuations (21) (c) versus 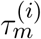, the excitatory (inhibitory) population are denoted by red (blue) circles. All the data here reported refer to MF simulations. The parameters are 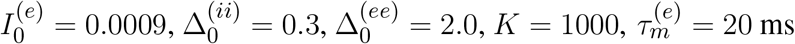.

#### 3.2.2 From Fluctuation Driven to Abnormally Synchronized Oscillations

As a second range of parameters, we consider the cut in the parameter plane shown in Fig. 1 (a) as a purple dashed line. For these parameters we report in Fig. 7 (a-b) the average in time of the excitatory and inhibitory population rate as a function of the excitatory DC current 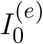. In particular, we compare network simulations (red and blue circles) with the MF results (red and blue lines). These predict a stable focus (solid lines) up to 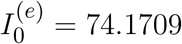, where a sub-critical Hopf bifurcation destabilizes such solution giving rise to an unstable focus (dashed lines). In panel (a) and (b) we have also reported as green dot-dashed lines the extrema of *R*^(*e*)^ and *R*^(*i*)^ corresponding to the unstable oscillations emerging at the Hopf bifurcation. We observe a good agreement for the time averaged activity with the MF results for currents smaller than that of the Hopf bifurcation, above which the MF model predicts a diverging solution.

**Figure 7.**
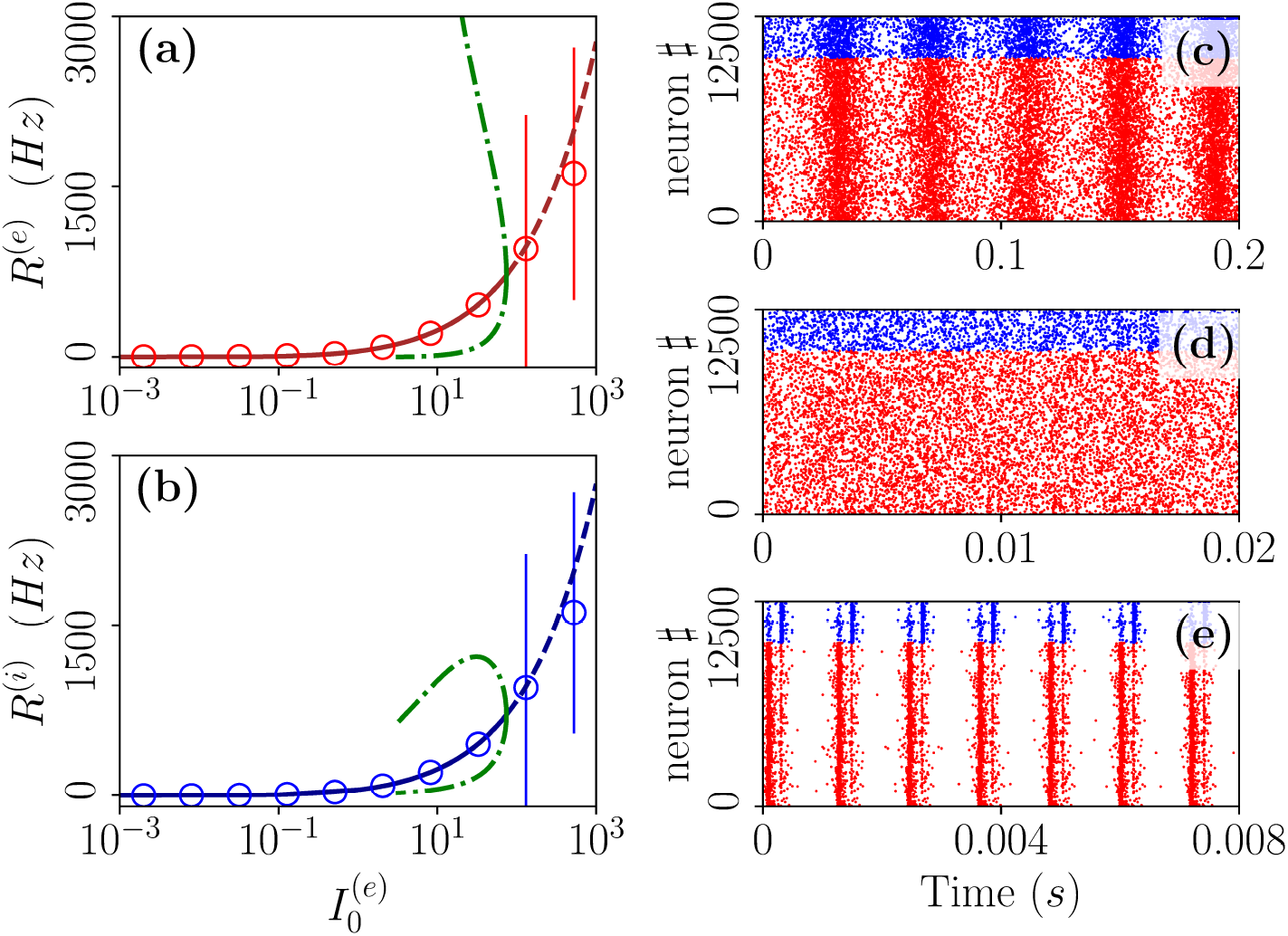
From fluctuation driven to abnormally synchronized oscillations. Firing rates *R*^(*e*)^ (a) and *R*^(*i*)^ (b) as a function of 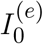 for E-I networks (circles) and neural mass model (lines) for the parameter cut corresponding to the dashed purple line in Fig. 1 (a). For the neural mass model: solid (dashed) line shows stable (unstable) focus solution 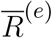 and 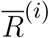; green dot-dashed lines refer to the extrema of *R*^(*e*)^(*R*^(*i*)^) for the unstable limit cycle present in region (II). The unstable limit cycle emerges at the sub-critical Hopf bifurcation for 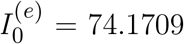 separating region (II) from (I), where the focus becomes unstable. Raster plots are reported for specific cases: namely, 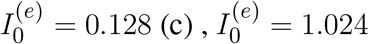 (d) and 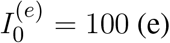 (e).Parameters as in Fig. 1, other parameters are set as 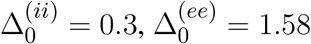, *K* = 1000, *N* ^(*e*)^ = 10000 and *N* ^(*i*)^ = 2500.

As reported in (Montbrió et al., 2015) when the network dynamics becomes strongly synchronous (as expected for very high common excitatory DC external current) the MF formulation fails since the population rates predicted within the MF formulation diverge. However, as shown in Fig. 7 (e) due to finite size effects we observe in the network a strongly synchronous COs of type O_S_ corresponding to the MF region (I) where the MF model predicts no stable solution. These abnormally synchronized oscillations are also characterized by a quite fast frequency of oscillation *ν*_CO_ *≃* 800 *−* 1000 Hz. Furthermore, similarly to the O_P_ oscillations they emerge due to a PING-like mechanism. This is evident from the raster plot in Fig. 7 (e), where excitatory neurons fire almost synchronously followed, after an extremely short delay, by the inhibitory ones whose activity silence all the network until the next excitatory burst. Quite astonishingly the mean population rates measured in the network are reasonably well captured by the MF solutions associated to the unstable focus even beyond the Hopf bifurcation, despite the network is now displaying COs (see Fig. 7 (a-b)).

Below the Hopf bifurcation, while the MF predicts only the existence of a stable focus the network dynamics reveals quite interesting features. As shown in Fig. 7 (d) the system dynamics is indeed asynchronous for intermediate current values, here 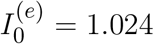, however at lower currents we observe fluctuation driven oscillations O_F_ as shown in Fig. 7 (c) for 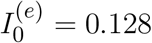. We will now investigate in more details the origin of these COs.

The emergence of COs in the network can be characterized in terms of the coherence indicator *ρ* (4) for the whole population of neurons. This indicator is reported in Fig. 8 (a) as a function of 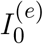 for the same parameters previously discussed in Fig. 7 and for two different values of the median in-degree : *K* = 100 (red circles) and *K* = 4000 (blue circles). For both values of *K*, we observe an almost discontinuous transition in the value of the coherence indicator at the sub-critical Hopf bifurcation from 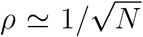,expected for an asynchronous dynamics, to values *ρ ≃* 1 corresponding to fully synchronization. This discontinuous transition leads to the emergence of abnormally synchronized oscillations O_S_ in the network. Moreover, at sufficiently high in-degrees we observe the emergence of a new coherent state for low DC currents 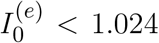 characterized by a finite value of the coherence indicator, namely *ρ ≃* 0.3. The origin of these oscillations can be better understood by examining the coefficient of variation *CV* averaged over the whole population, this is reported in Fig. 8 (c) for the same interval of excitatory DC current and the same in-degrees as in Fig. 8 (a). It is evident that the *CV* assumes finite values only for small input currents, namely 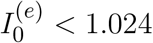, indicating the presence of not negligible fluctuations in the network dynamics. Furthermore, by increasing *K* these fluctuations, as measured by the *CV*, increases as expected for a balanced network. This analysis suggests that these oscillations cannot exist in absence of fluctuations in the network and therefore they are of the O_F_ type. Furthermore, the network should be sufficiently connected in order to sustain these COs, as one can understand from Fig. 8 (b) and (d), where *ρ* and *CV* are reported as a function of *K* for three different values of 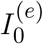. Indeed, for these parameter values no O_F_ oscillation is observable for *K <* 400, even in presence of finite values of the *CV*.

**Figure 8.**
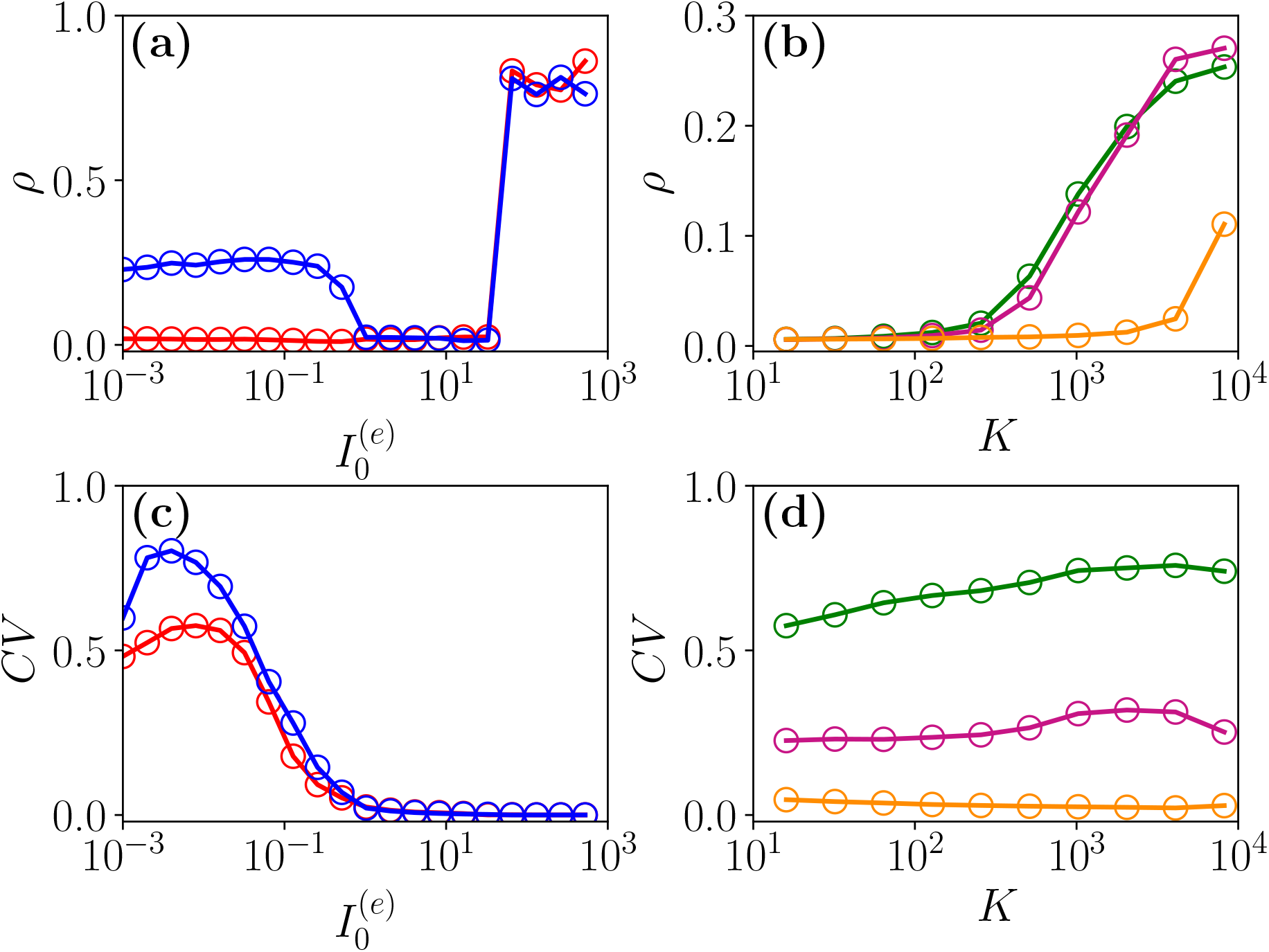
From fluctuation driven to abnormally synchronized oscillations. Coherence indicator *ρ* (4) for the whole network of excitatory and inhibitory neurons versus the excitatory DC current 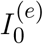 (a) and the median in-degree *K* (c). Coefficient of variation *CV* for the whole network versus 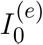 (b) and *K* (d). In panel (a) and (c) the symbols refer to different values of the median in-degree:namely, *K* = 100 (red circles) and *K* = 4000 (blue circles). In (b) and (d) the symbols refer to different excitatory DC currents: namely, 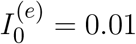 (green circles), 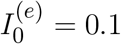 (purple circles) and 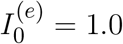 (orange circles). Parameters as in Fig. 1, other parameters 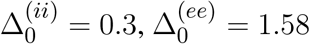, *N* ^(*e*)^ = 40000 and *N* ^(*i*)^ = 10000.

As previously discussed in (di Volo and Torcini, 2018), the balance between excitation and inhibition generates endogenous fluctuations that modifies the collective dynamics with respect to that predicted by the MF model, where the heterogeneity of the input currents, due to distributed in-degrees, is taken in account only as a quenched form of disorder and not as a dynamical source of noise. However, also from this simplified MF formulation one can obtain relevant information on the O_F_ oscillations, indeed as we will see in the next sub-section the relaxation frequencies towards the stable MF focus represent a good estimation of the oscillation frequencies measured in the network. This suggests that the fluctuations present at the network level can sustain COs by continuously exciting the focus observed in the effective MF model with quenched disorder.

#### 3.2.3 Fluctuation driven oscillations: from quasi-periodicity to frequency locking

As announced, this sub-section will be devoted to the characterization of the fluctuation driven oscillations O_*F*_ emerging in region (II) reported in Fig. 1. As the MF is now characterized by a stable focus with two couples of complex conjugate eigenvalues there are two frequencies that can be excited by neurons’ irregular firing. Accordingly, as reported in (di Volo and Torcini, 2018), we expect the collective dynamics to be characterized by a quasi-periodic dynamics with two (incommensurable) frequencies. These frequencies can be estimated by computing the power spectrum *S*(*ν*) of global quantities, e.g. mean membrane potential *V* (*t*). In the case of a periodic dynamics *S*(*ν*) is characterized by one main peak in correspondence of the CO frequency and minor peaks at its harmonics, while in the quasi-periodic case the power spectrum shows peaks located at the two fundamental frequencies and at all their linear combinations. Indeed, as shown in Fig. 9 (a) the power spectrum exhibit several peaks over a continuous profile and the peak frequencies can be obtained as a linear combination of two fundamental frequencies (*ν*_1_, *ν*_2_). As already mentioned, the noisy background is due to the fluctuations present in the balanced network. It is evident from Fig. 9 (b) that these two fundamental frequencies are well reproduced by the two relaxation frequencies 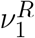 and 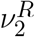 towards the MF focus, in particular for 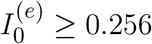. At smaller currents, while the first frequency is well reproduced by 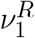, the second one is under-estimated by 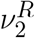. This is due to the phenomenon of frequency locking among the two collective rhythms present in the system: when the two frequencies become commensurable we observe a common periodic CO. The locking order can be estimated by plotting the ratio between the two frequencies, indeed for low currents and *K* = 8192 the ratio is almost constant and equal to four denoting a 1:4 frequency locking (see Fig. 9 (c)). Furthermore, by fixing 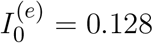 and by varying *K* the ratio *ν*_1_*/ν*_2_ can display different locked states, passing from a locking of type 1 : 2 at low *K* to 1 : 4 at larger values, as shown in the inset of Fig. 9 (c).

**Figure 9.**
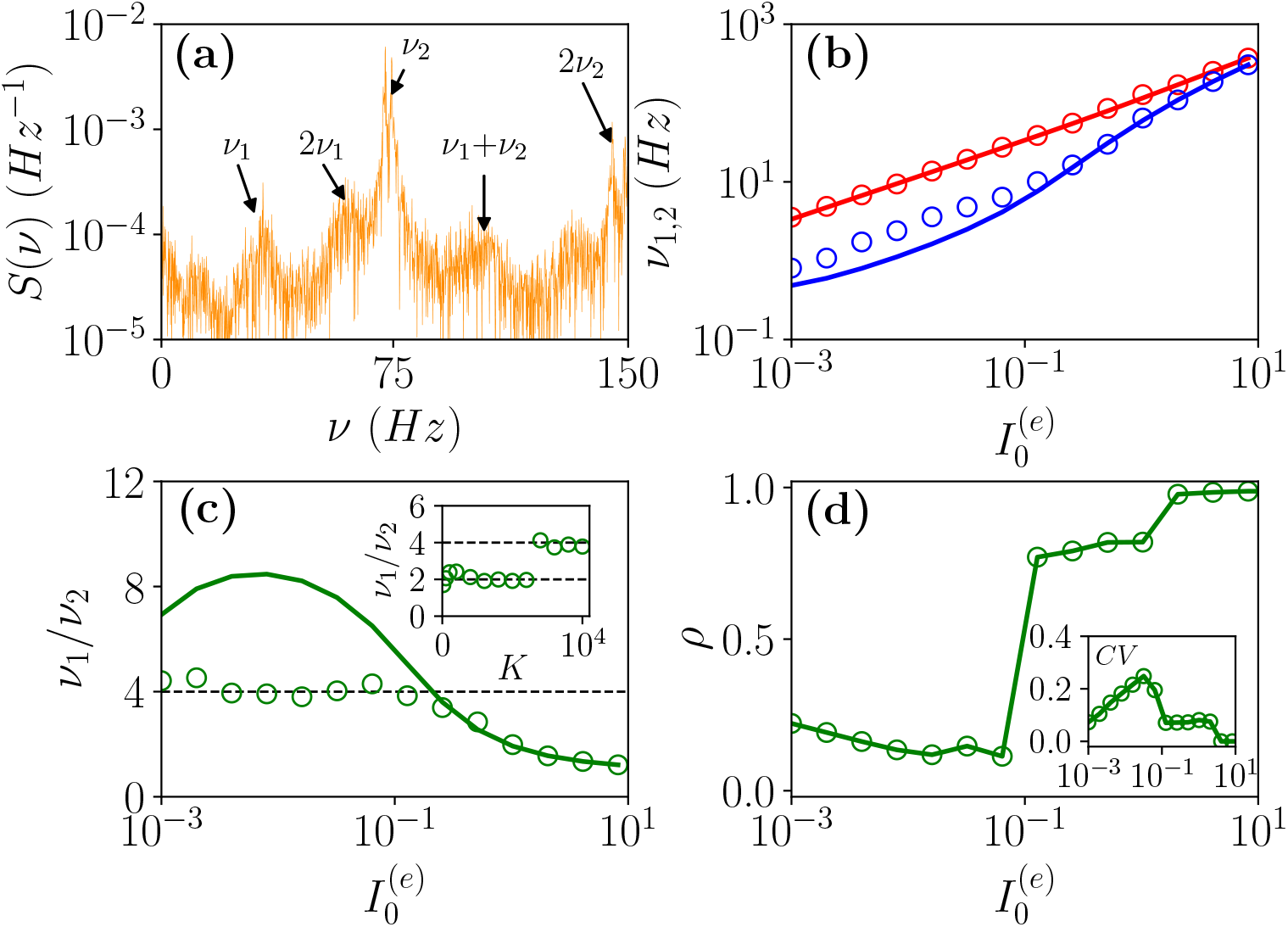
From quasi-periodicity to frequency locking. (a) Power spectra *S*(*ν*) of the mean membrane potential obtained from network simulations. (b) The two fundamental frequencies *ν*_1_(*ν*_2_) versus 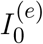. (c) Frequency ratio *ν*_1_*/ν*_2_ versus 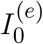, in the inset *ν*_1_*/ν*_2_ is shown versus *K*. (d) Coherence parameter *ρ* versus 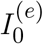, in the inset the corresponding *CV* is reported. In (b-c) the symbols (solid lines) refer to *ν*_1_ and *ν*_2_ as obtained from the peaks of the power spectra *S*(*ν*) for *V* (*t*) obtained from the network dynamics (to the the two relaxation frequencies 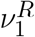 and 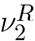 associated to the stable focus solution for the MF). Parameters as in Fig. 1, other parameters are set as 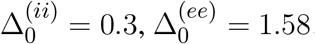, *N* ^(*e*)^ = 80000, *N* ^(*i*)^ = 20000, *K* = 8192 and 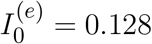 in the inset of panel (c).

As evident from Fig. 9 (b) and (c), the locking phenomenon arises only in the network simulations and it is not captured by the MF model. Furthermore, frequency locking occurs at low currents 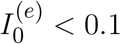 where the dynamics of the neurons is driven by the intrinsic current fluctuations present in the network, but not in the MF. Indeed for low DC currents the level of synchronization within the populations measured by *ρ* decreases with 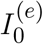, while the *CV* increases (as shown in Fig. 8 (d)). These features suggest that this phenomenon is somehow similar to what reported in (Meng and Riecke, 2018) for two coupled inhibitory neural populations subject to external uncorrelated noise. The authors in (Meng and Riecke, 2018) observed an increase of the locking region among collective rhythms by increasing the amplitude of the additive noise terms, this joined to a counter-intuitive decrease of the level of synchronization among the neurons within each population. However, in (Meng and Riecke, 2018) the neurons are subject to independent external noise sources, while in our case the sources of fluctuations are intrinsic to the system and induced by the structural heterogeneity. Due to the network sparseness the current fluctuations experienced by each neuron can be assumed to be indeed uncorrelated (Brunel and Hakim, 1999). Therefore we are facing a new phenomenon that we can identify as a frequency locking of collective rhythms promoted by self-induced uncorrelated fluctuations. Indeed, the locking disappears for increasing external DC currents 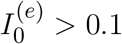, when the coherence parameter *ρ* displays an abrupt jump towards higher values and the *CV ≃* 0, thus indicating that in this regime the neuron dynamics becomes essentially mean driven.

#### 3.2.4 COs’ amplitude and frequency dependence on the in-degree and DC current

The dynamics of balanced networks is usually characterized in the limit *N >> K >>* 1 by the emergence of a self-sustained asynchronous regime. However, limit cycle solutions have been already reported for balanced networks in the seminal paper by Van Vreeswijk and Sompolinsky (van Vreeswijk and Sompolinsky, 1996). These solutions can be either unbalanced or balanced, however in this latter case they were characterized by vanishing small oscillations’ amplitudes, scaling as *K*^*−*1*/*2^. As a matter of fact, the authors in (van Vreeswijk and Sompolinsky, 1996) have shown that balanced COs are not observable in their model in the limit *N >> K→ ∞*, but only for finite *K*. Therefore, it is important to address also in our case if COs can still be observable in the limit *N >> K >>* 1 and which are their characteristics. Thus, in the following we will investigate the behaviour of the frequencies and amplitudes of COs for diverging *K* and for increasing external DC currents.

Let us first consider fluctuation driven O_*F*_ oscillations, in this case we have an analytical prediction (20) for the scaling of the fundamental frequencies 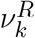 associated to the relaxation towards the macroscopic focus, which should grow proportionally to 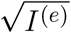. As shown in Figs. 10 (a) and (c), indeed this scaling is clearly observable for sufficiently large *K* and 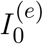. It is also evident the extreme good agreement between results obtained from the network simulations and the theoretical predictions (20), at least for the values of *K* reachable with our simulations. Furthermore, the COs’ frequencies cover an extremely large range of values from few Hz to KHz and this range of frequencies can be spanned by varying *K* and the external DC current 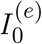 as shown in Fig. 10 (a) and (c).

**Figure 10.**
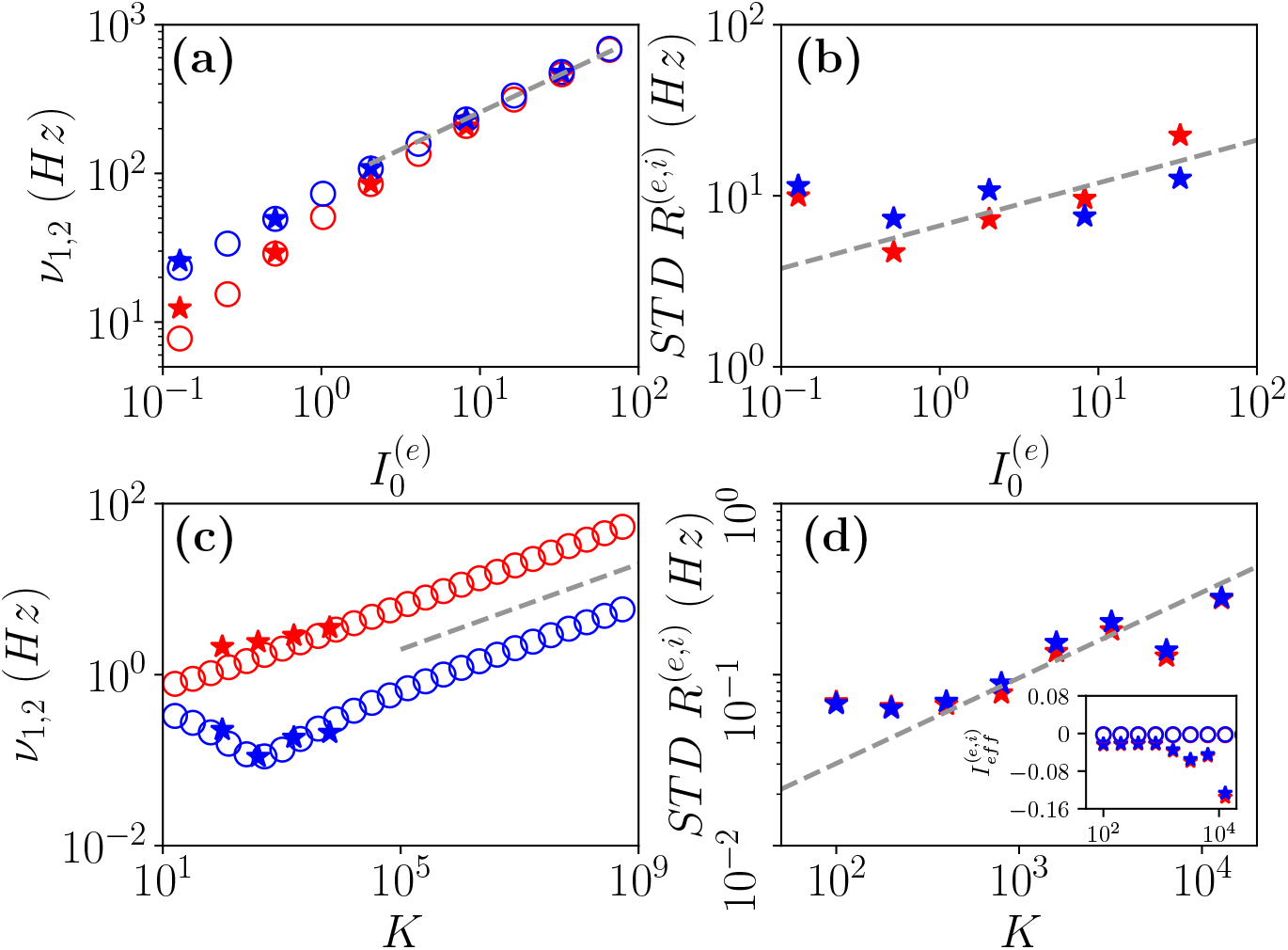
Frequencies and amplitudes of O_*F*_ oscillations. The two fundamental frequencies *ν*_1_ and *ν*_2_ versus 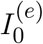 (a) and *K* (c) for network simulations (stars) and for the MF (circles) (the latter ones are the two relaxation frequencies 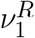 and 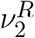 associated to the stable focus). Standard deviations of the firing rates for the excitatory (red) and inhibitory (blue) populations versus versus 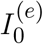 (b) and *K* (d) as obtained from network simulations. In the inset in (d) the effective mean input currents 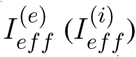 of the excitatory (inhibitory) population are shown versus *K* for the MF focus (circles) and as estimated from the network dynamics (stars). The dashed line in panel (a) (panel (c)) corresponds to a power law-scaling 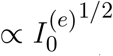 for the frequencies of the COs and in panel (b) (panel (d)) to a power law-scaling 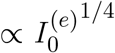 (∞ *K*^1/2^) for their amplitudes. Parameters as in Fig. 1, other parameters: (a-b) *K* = 1000, 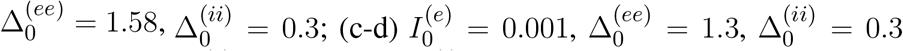; for the network simulations we employed *N* ^(*e*)^ = 80000 and *N* ^(*i*)^ = 20000.

The amplitudes of the COs have been measured by estimating the standard deviation of the population firing rate for the excitatory (inhibitory) population STD *R*^(*e*)^ (STD *R*^(*i*)^). These quantities are displayed for O_*F*_ oscillations in Figs. 10 (b) and (d) as a function of 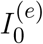 and *K*, respectively. Clearly for the estimation of these quantities we can rely only on network simulations, the numerical results suggest a divergence of the amplitudes of the O_F_ oscillations with *K* with a power-law increase *∝K*^1*/*2^ for *K >* 300, while the dependence on 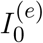 suggests a limited increase consistent with a scaling 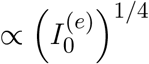 over almost two decades.

These results seem to indicate that for *N >> K → ∞* the network will display fluctuation driven oscillations of wider and wider amplitudes, joined to an increase in the level of synchronization, as confirmed by direct measurements of the indicator *ρ* (not shown here). This behaviour is associated to a reduction of the periods of such oscillations with *K*, thus for sufficiently large in-degrees we expect to converge towards a sort of ictal state characterized by abnormally fast and synchronized oscillations. However, at variance with the ictal state previously analyzed, that was characterized by an extremely regular dynamics of the neurons (see Fig. 8), here the system always exhibits a fluctuation driven dynamics, since we measured *CV ≃* 0.6 *−* 0.8 at least in the range *K ≃* 100 *−* 10^4^ accessible to network simulations. This is confirmed by the analysis of the mean effective input currents 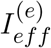 and 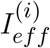 shown in the inset of Figs. 10 (d). For the MF focus the dynamics appear as almost exactly balanced for all the considered median in-degree *K* since 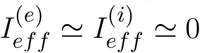. These results are confirmed by the network simulations for *K ≤* 1000. However, at larger *K* the network dynamics is on average dominated by the inhibitory drive and therefore the neurons should be on average slightly sub-threshold. This does not prevent the emergence of COs driven by fluctuations at large *K*, as indeed observed.

Analogous scalings for the frequencies and amplitudes of fluctuation driven COs have been recently found for a purely inhibitory balanced network (Segneri et al., 2021), thus confirming the generality of this scenario, at least for QIF neurons. The only difference being in the value of the exponent controlling the power-law increase of the COs’ amplitudes with 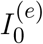.

Let us now examine frequencies and amplitudes of O_*P*_ oscillations. As shown in Figs. 11 (a) and (c), the frequencies *ν*_*CO*_ as estimated from the MF model (open circles) reveal an almost perfect increase proportional to 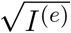 analogous to the one reported for O_*F*_ oscillations. The data obtained from network simulations (stars) converge towards the MF results for sufficiently large *K* and 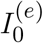.

**Figure 11.**
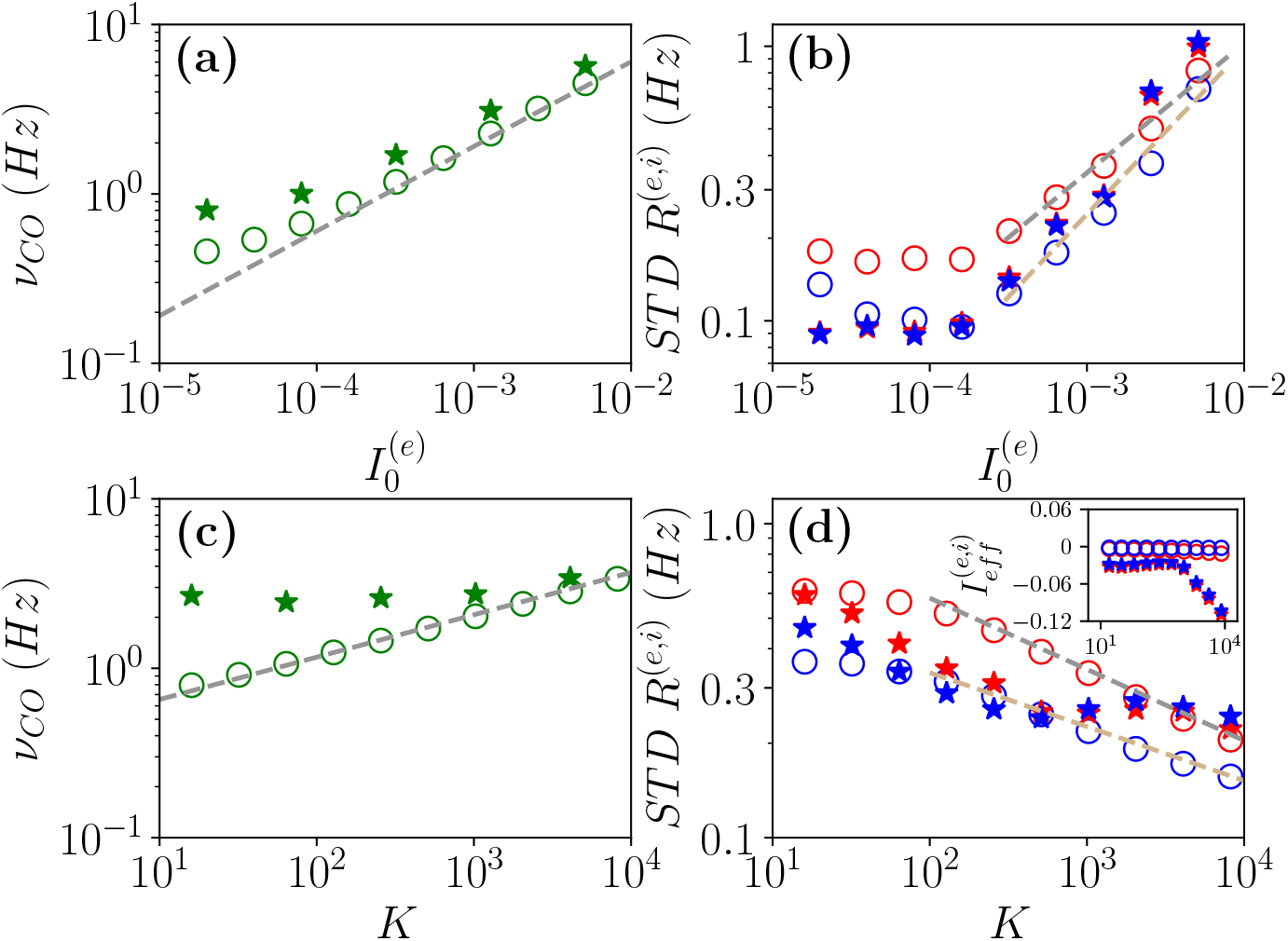
Frequencies and amplitudes of O_*P*_ oscillations. COs’ frequency *ν*_*CO*_ versus 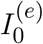 (a) and *K* (c) and standard deviations of the firing rates for the excitatory (red) and inhibitory (blue) populations versus versus 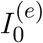 (b) and *K* (d). The data obtained from network (MF) simulations are denoted as stars (circles). The dashed line in panel (a) (panel (c)) corresponds to a power law-scaling 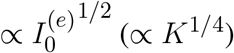 for the frequencies. In the inset in (d) the effective mean input currents 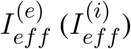 of the excitatory (inhibitory) population are shown versus *K* for the MF (circles) and as estimated from the network dynamics (stars). The dashed lines in panel (b) refer to a fitting to the MF data with a power law 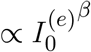, where *β* ≃ 0.47(*β ≃* 0.60) for the excitatory (inhibitory) population; those in panel (d) to a fitting to the MF data with a power law *∝K*^*−γ*^, where *γ ≃*0.23 (*γ≃* 0.17) for the excitatory (inhibitory) population. The data reported in (a-b) and (c-d) refer to the open circles in Fig. 1 (a) and (b), respectively. For network simulation we employed *N* ^(*e*)^ = 80000 and *N* ^(*i*)^ = 20000.

The amplitudes of these COs reveal power-law increases with 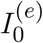 for sufficiently large DC currents, as shown in Figs. 11 (b), furthermore the MF results and the network simulations show a reasonable agreement over the examined range. This behaviour is analogous to O_*F*_ oscillations. The main difference with respect to fluctuation driven oscillations, is the dependence of the COs’ amplitudes on the median in-degree *K*. As evident from Figs. 11 (d), the MF results (open circles) for *K >* 100 tends to vanish as *∝K*^*−γ*^ with *γ ≃* 0.2. However, network simulations show a saturation of such amplitudes for *K >* 1000. At the same time, the MF is perfectly balanced in the whole range of examined in-degrees, since 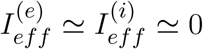, while the network simulations reveal balanced effective input currents up to *K ≃* 1000 and above such median in-degree a prevalence of the inhibitory drive (see inset of Figs. 11 (d)). The origin of these discrepancies between the MF and the network simulations at large *K* is unclear and deserves future investigations. However, the MF results indicate that these balanced COs in the limit *N >> K >>* 1 exhibit vanishing small amplitudes joined to higher and higher frequency, thus suggesting that the asymptotic state would probably be a balanced asynchronous regime.

## 4 CONCLUSIONS

We have characterized the dynamics of a sparse balanced E-I QIF network with Lorentzian distributed in-degrees. This peculiar choice of the distribution has allowed us to derived an exact low dimensional neural mass model describing the MF dynamics of the network in terms of the mean membrane potentials and of the population rates of the two populations (Montbrió et al., 2015; di Volo and Torcini, 2018). The low-dimensionality of the MF equations enabled us to study analytically the stationary solutions and their stability for any finite value of the median in-degree *K*, as well as to obtain the bifurcation diagrams associated to the model and to identify the possible macroscopic states.

The stationary solutions of the MF correspond to the asynchronous regime, which is the regime usually analyzed in the context of balanced dynamics (van Vreeswijk and Sompolinsky, 1996; Renart et al., 2010; Litwin-Kumar and Doiron, 2012). In the present case we have analytically obtained the stationary solutions for the mean membrane potentials and average firing rates for Lorentzian distributed in-degrees for any finite value of the median *K* and for a HWHM scaling as 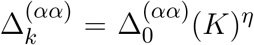 with *η* = 1*/*2. The MF estimations for the population firing rates are pretty well reproduced by the network simulations in the examined range of in-degrees *K*. Furthermore, from the analytic expression of the stationary firing rates (8) it is evident that for *K >>* 1 the asymptotic rates would not depend on the structural heterogeneity and correspond to those usually found for balanced homogeneous or Erdös-Renyi networks (van Vreeswijk and Sompolinsky, 1996; Monteforte and Wolf, 2010). This is due to the fact that the ratio 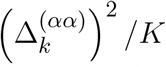 remains constant for *K → ∞*. The final scenario will depend on the scaling exponent *η*, in particular by assuming *η* = 3*/*4 the asymptotic firing rates 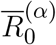 will explicitly depend on the parameters 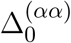 controlling the structural heterogeneity. Whenever *η >* 3*/*4 the balanced state breaks down and we face a situation similar to those investigated in (Landau et al., 2016; Pyle and Rosenbaum, 2016), therefore balance can be recovered either rewiring the post-synaptic connections (Pyle and Rosenbaum, 2016) or by introducing some sort of homeostatic plasticity or of spike-frequency adaptation (Landau et al., 2016).

However, despite the system approaches a balanced state, as testified by the fact that the effective input currents converge to finite values 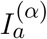 for *K → ∞*, the balanced regime is not necessarily a sub-threshold one. Indeed, we have observed that we can have either sub-threshold or supra-threshold situations depending on the model parameters in agreement with the results previously reported in (Lerchner et al., 2006). Moreover, the excitatory and inhibitory populations can achieve balanced regimes characterized by different asymptotic dynamics, i.e. with 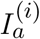 and 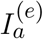 of opposite sign.

While at a macroscopic level the population activity for *N >> K >>* 1 approaches essentially that of a homogeneous balanced system, as shown in Fig. 2 (a) and (b), the structural heterogeneity has a large influence on the single neuron dynamics, at least at finite *K* and finite investigation times. In particular, in analogy with experiments (Gentet et al., 2010; Mongillo et al., 2018) we considered a situation where the inhibitory drive prevails on the excitatory one. In this condition microscopically the neural populations splits in three groups: silent neurons, definitely sub-threshold; bulk neurons, which are fluctuation driven; and mean driven outlier neurons. In particular, excitatory (inhibitory) neurons with low (high) intra-population in-degrees are silenced due to the prevalence of synaptic inhibition. The silent neurons represent 6-10 % of the whole population in agreement with experimental results for the mice cortex (O’Connor et al., 2010). Bulk neurons have in-degrees in proximity of the median and their firing rates approach the MF solution 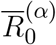 for increasing *K*, Outlier neurons represent a definitely minority group almost disconnected from their own population, whose asymptotic behaviour for *K >>* 1 is controlled by the sign of the effective mean input current.

The emergence of COs is observable in this balanced network whenever the level of heterogeneity in the inhibitory population is not too large, thus suggesting that the coherence among inhibitory neurons is fundamental to support collective rhythms (Whittington et al., 2000). Indeed we observed two main mechanisms leading to COs: one that can be identified as PING-like and another one as fluctuation driven. The PING-like mechanism is present whenever the excitatory neurons are able to deliver an almost synchronous excitatory volley that in turn elicits a delayed inhibitory one. The period of the COs is determined by the recovery time of the excitatory neurons from the stimulus received from the inhibitory population. This mechanism is characterized by a delay between the firing of the pyramidal cells and the interneuronal burst as reported also in many experiments (Buzsáki and Wang, 2012). We have shown that this delay tends to vanish when the inhibitory action increases leading the system from a balanced situation to a definitely sub-threshold condition where the neural activity is completely controlled by fluctuations. In this latter case the excitatory and inhibitory neurons fire almost simultaneously driven by the current fluctuations. These transform the relaxation dynamics towards a stable focus, observable in the MF, to sustained COs via a mechanism previously reported for inhibitory networks (di Volo and Torcini, 2018; Bi et al., 2020).

The PING-like COs undergo period doubling cascades by varying *K* and/or 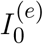 finally leading to collective chaos (Nakagawa and Kuramoto, 1993; Shibata and Kaneko, 1998). The nature of this chaotic behaviour is definitely macroscopic since it is captured by the neural mass model obtained within the MF formulation, as shown by analysing the corresponding Lyapunov spectrum. This kind of chaos implies irregular temporal fluctuations joined to a coherence at the spatial level over a large part of the network resembling coherent fluctuations observed across spatial scales in the neocortex (Volgushev et al., 2011; Smith and Kohn, 2008; Okun et al., 2012; Achermann et al., 2016). Collective or coherent chaos has been previously shown to be a ubiquitous feature for balanced random spiking neural networks massively coupled, where *K* is proportional to *N* (Ullner et al., 2018; Politi et al., 2018). *Here, we have generalized such result to balanced random networks with sparse connectivity, where K* is independent by *N*. Recently, it has been claimed that the presence of a structured feed forward connectivity in a random network is needed to observe coherent chaos (Landau and Sompolinsky, 2018). However, as evident from our results and those reported in (Ullner et al., 2018; Politi et al., 2018) coherent chaos can naturally emerge in a recurrent neural network in absence of any structured connectivity introduced *ad hoc* to promote collective behaviours. Furthermore, we have shown that collective chaos can emerge in random balanced networks with instantaneous synapses and in absence of any delay, see also (Ullner et al., 2018). This seems to contrast with what happens in globally coupled neural systems, where collective chaos has been so far reported in presence of some extrinsic time scale, which can be a synaptic time scale, as in (Olmi et al., 2011) for two coupled excitatory populations of LIF neurons and in (Ceni et al., 2020) for two coupled inhibitory QIF populations, or a time delay as shown in (Luccioli and Politi, 2010; Pazó and Montbrió, 2016; Devalle et al., 2018) for single inhibitory LIF and QIF populations.

Fluctuation driven COs are usually observable in our system as quasi-periodic collective motions characterized by two incommensurate frequencies. However, whenever the current fluctuations become sufficiently strong the two frequencies can lock and give rise to a collective periodic motion. Furthermore, the locking region is characterized by a low level of synchrony in the network. These results resemble those reported in (Meng and Riecke, 2018) for two interconnected inhibitory neural networks subject to external uncorrelated noise. In particular, the authors have shown that uncorrelated noise sources enhance synchronization and frequency locking among the COs displayed by the two networks, despite the noise reduces the synchrony among neurons within each network. At variance with (Meng and Riecke, 2018), in our case the noise sources are intrinsic to the neural dynamics, but they can be as well considered as uncorrelated due to the sparseness in the connections (Brunel and Hakim, 1999; Brunel, 2000). Therefore we are reporting a new example of frequency locking among collective rhythms promoted by self-induced uncorrelated fluctuations.

The frequencies of COs grows proportionally to the square root of the external excitatory DC current, as suggested by analytical arguments and confirmed by numerical simulations. This on one side allows, simply by varying the parameters 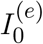 or *K*, to cover with our model a broad range of COs’ frequencies analogous to those found experimentally in the cortex (Chen et al., 2017). On another side it implies that the frequencies of COs diverge as *K*^1*/*4^. This result joined to the fact that the amplitudes of the fluctuation driven (PING-like) COs tend to vanish (to diverge) for increasing *K* suggests that, despite balanced COs are observable for any finite *K* in our model, in the limit *N >> K >>* 1 the model could display only two coexisting regimes. These are an asynchronous balanced regime and a fully synchronized one characterized by infinitely fast population bursts.

As a last issue, we would like to remark that our neural mass model has been derived by taking into account the random fluctuations due to the sparseness in the network connectivity only as a quenched disorder affecting the distribution of the effective synaptic couplings (Montbrió et al., 2015; di Volo and Torcini, 2018). The current fluctuations can be correctly incorporated in a MF formulation by developing a Fokker-Planck formalism for the problem, however this will give rise to high (infinite) dimensional MF models (Brunel and Hakim, 1999; Brunel, 2000). We are currently working on developing reduction formalisms for the Fokker-Planck equation to obtain low dimensional neural mass models for the evolution of the mean membrane potential and of the population rate which will include the intrinsic current fluctuations as a Poissonian term (Goldobin et al., 2021; Segneri et al., 2021). A further improvement would consist in including in a self-consistent manner the current fluctuations in the MF formulation for balanced sparse networks. This has been done for homogeneous (Ullner et al., 2020) and heterogeneous (Lerchner et al., 2006) LIF neurons. The self-consistent MF results have been obtained via refined iterative procedures based on the assumption that the synaptic input currents are the superposition of independent single neuron dynamics. However, such approaches neglect the correlations among the neurons and it is unclear how they can be extended to system displaying COs.

## AUTHOR CONTRIBUTIONS

HB and MdV performed the simulations and data analysis. MdV and AT were responsible for the state-of-the-art review and the paper write-up. All the authors conceived and planned the research.

## ACKNOWLEDGMENTS

The authors acknowledge useful extremely useful discussion with D.G. Goldobin, G. Mongillo, E. Montbrió, S. Olmi, and A. Politi.

## DATA AVAILABILITY STATEMENT

The numerical programs and datasets for this study are availabl upon request.

## REFERENCES

Achermann, P., Rusterholz, T., Dürr, R., König, T., and Tarokh, L. (2016). Global field synchronization reveals rapid eye movement sleep as most synchronized brain state in the human eeg. Royal Society open science 3, 160201

Angulo-Garcia, D., Luccioli, S., Olmi, S., and Torcini, A. (2017). Death and rebirth of neural activity in sparse inhibitory networks. New Journal of Physics 19, 053011

Barral, J. and Reyes, A. D. (2016). Synaptic scaling rule preserves excitatory–inhibitory balance and salient neuronal network dynamics. Nature neuroscience 19, 1690

Benettin, G., Galgani, L., Giorgilli, A., and Strelcyn, J.-M. (1980). Lyapunov characteristic exponents for smooth dynamical systems and for hamiltonian systems; a method for computing all of them. part 1: Theory. Meccanica 15, 9–20

Bi, H., Segneri, M., di Volo, M., and Torcini, A. (2020). Coexistence of fast and slow gamma oscillations in one population of inhibitory spiking neurons. Physical Review Research 2, 013042

Brunel, N. (2000). Dynamics of sparsely connected networks of excitatory and inhibitory spiking neurons. Journal of Computational Neuroscience 8, 183–208. doi:10.1023/A:1008925309027

Brunel, N. and Hakim, V. (1999). Fast global oscillations in networks of integrate-and-fire neurons with low firing rates. Neural computation 11, 1621–1671

Bruno, R. M. and Sakmann, B. (2006). Cortex is driven by weak but synchronously active thalamocortical synapses. Science 312, 1622–1627

Buzsáki, G. and Wang, X.-J. (2012). Mechanisms of gamma oscillations. Annual review of neuroscience 35, 203–225

Ceni, A., Olmi, S., Torcini, A., and Angulo-Garcia, D. (2020). Cross frequency coupling in next generation inhibitory neural mass models. Chaos: An Interdisciplinary Journal of Nonlinear Science 30, 053121

Chen, G., Zhang, Y., Li, X., Zhao, X., Ye, Q., Lin, Y., et al. (2017). Distinct inhibitory circuits orchestrate cortical beta and gamma band oscillations. Neuron 96, 1403–1418

Dehghani, N., Peyrache, A., Telenczuk, B., Le Van Quyen, M., Halgren, E., Cash, S. S., et al. (2016). Dynamic balance of excitation and inhibition in human and monkey neocortex. Scientific reports 6, 23176

Destexhe, A. and Paré, D. (1999). Impact of network activity on the integrative properties of neocortical pyramidal neurons in vivo. Journal of neurophysiology 81, 1531–1547

Devalle, F., Montbrió, E., and Pazó, D. (2018). Dynamics of a large system of spiking neurons with synaptic delay. Physical Review E 98, 042214

di Volo, M. and Torcini, A. (2018). Transition from asynchronous to oscillatory dynamics in balanced spiking networks with instantaneous synapses. Phys. Rev. Lett. 121, 128301

Ermentrout, B. (2007). XPPAUT. Scholarpedia 2, 1399. doi:10.4249/scholarpedia.1399. Revision #136177

Ermentrout, G. B. and Kopell, N. (1986). Parabolic bursting in an excitable system coupled with a slow oscillation. SIAM Journal on Applied Mathematics 46, 233–253

Gentet, L. J., Avermann, M., Matyas, F., Staiger, J. F., and Petersen, C. C. (2010). Membrane potential dynamics of gabaergic neurons in the barrel cortex of behaving mice. Neuron 65, 422–435

Goldobin, D. S., Di Volo, M., and Torcini, A. (2021). A reduction methodology for fluctuation driven population dynamics. Phys. Rev. Lett.

Golomb, D. (2007). Neuronal synchrony measures. Scholarpedia 2, 1347. doi:10.4249/scholarpedia.1347

Haider, B., Duque, A., Hasenstaub, A. R., and McCormick, D. A. (2006). Neocortical network activity in vivo is generated through a dynamic balance of excitation and inhibition. Journal of Neuroscience 26, 4535–4545

Isaacson, J. S. and Scanziani, M. (2011). How inhibition shapes cortical activity. Neuron 72, 231–243

Kadmon, J. and Sompolinsky, H. (2015). Transition to chaos in random neuronal networks. Phys. Rev. X 5, 041030. doi:10.1103/PhysRevX.5.041030

Landau, I. D., Egger, R., Dercksen, V. J., Oberlaender, M., and Sompolinsky, H. (2016). The impact of structural heterogeneity on excitation-inhibition balance in cortical networks. Neuron 92, 1106–1121

Landau, I. D. and Sompolinsky, H. (2018). Coherent chaos in a recurrent neural network with structured connectivity. PLoS computational biology 14, e1006309

Le Van Quyen, M., Muller, L. E., Telenczuk, B., Halgren, E., Cash, S., Hatsopoulos, N. G., et al. (2016). High-frequency oscillations in human and monkey neocortex during the wake–sleep cycle. Proceedings of the National Academy of Sciences 113, 9363–9368

Lefort, S., Tomm, C., Sarria, J.-C. F., and Petersen, C. C. (2009). The excitatory neuronal network of the {C2} barrel column in mouse primary somatosensory cortex. Neuron 61, 301 – 316. doi:https://doi.org/10.1016/j.neuron.2008.12.020

Lehnertz, K., Bialonski, S., Horstmann, M.-T., Krug, D., Rothkegel, A., Staniek, M., et al. (2009). Synchronization phenomena in human epileptic brain networks. Journal of neuroscience methods 183, 42–48

Lerchner, A., Ursta, C., Hertz, J., Ahmadi, M., Ruffiot, P., and Enemark, S. (2006). Response variability in balanced cortical networks. Neural computation 18, 634–659

Litwin-Kumar, A. and Doiron, B. (2012). Slow dynamics and high variability in balanced cortical networks with clustered connections. Nat Neurosci 15, 1498–1505

Luccioli, S. and Politi, A. (2010). Irregular collective behavior of heterogeneous neural networks. Phys. Rev. Lett. 105, 158104. doi:10.1103/PhysRevLett.105.158104

Meng, J. H. and Riecke, H. (2018). Synchronization by uncorrelated noise: interacting rhythms in interconnected oscillator networks. Scientific reports 8, 1–14

Mongillo, G., Rumpel, S., and Loewenstein, Y. (2018). Inhibitory connectivity defines the realm of excitatory plasticity. Nature neuroscience 21, 1463–1470

Montbrió, E., Pazó, D., and Roxin, A. (2015). Macroscopic description for networks of spiking neurons. Physical Review X 5, 021028

Monteforte, M. and Wolf, F. (2010). Dynamical entropy production in spiking neuron networks in the balanced state. Phys. Rev. Lett. 105, 268104. doi:10.1103/PhysRevLett.105.268104

Nakagawa, N. and Kuramoto, Y. (1993). Collective chaos in a population of globally coupled oscillators. Progress of Theoretical Physics 89, 313–323

O’Connor, D. H., Peron, S. P., Huber, D., and Svoboda, K. (2010). Neural activity in barrel cortex underlying vibrissa-based object localization in mice. Neuron 67, 1048–1061

Okun, M. and Lampl, I. (2008). Instantaneous correlation of excitation and inhibition during ongoing and sensory-evoked activities. Nature neuroscience 11, 535

Okun, M., Yger, P., Marguet, S. L., Gerard-Mercier, F., Benucci, A., Katzner, S., et al. (2012). Population rate dynamics and multineuron firing patterns in sensory cortex. Journal of Neuroscience 32, 17108– 17119

Olmi, S., Politi, A., and Torcini, A. (2011). Collective chaos in pulse-coupled neural networks. EPL (Europhysics Letters) 92, 60007

Ostojic, S. (2014). Two types of asynchronous activity in networks of excitatory and inhibitory spiking neurons. Nat Neurosci 17, 594–600

Ott, E. (2002). Chaos in dynamical systems (Cambridge university press)

Pazó, D. and Montbrió, E. (2016). From quasiperiodic partial synchronization to collective chaos in populations of inhibitory neurons with delay. Physical review letters 116, 238101

Pikovsky, A. and Politi, A. (2016). Lyapunov exponents: a tool to explore complex dynamics (Cambridge University Press)

Politi, A., Ullner, E., and Torcini, A. (2018). Collective irregular dynamics in balanced networks of leaky integrate-and-fire neurons. The European Physical Journal Special Topics 227, 1185–1204

Pyle, R. and Rosenbaum, R. (2016). Highly connected neurons spike less frequently in balanced networks. Physical Review E 93, 040302

Renart, A., de la Rocha, J., Bartho, P., Hollender, L., Parga, N., Reyes, A., et al. (2010). The asynchronous state in cortical circuits. Science 327, 587–590. doi:10.1126/science.1179850

Rosenbaum, R. and Doiron, B. (2014). Balanced networks of spiking neurons with spatially dependent recurrent connections. Physical Review X 4, 021039

Roxin, A., Brunel, N., Hansel, D., Mongillo, G., and van Vreeswijk, C. (2011). On the distribution of firing rates in networks of cortical neurons. Journal of Neuroscience 31, 16217–16226

Segneri, M., di Volo, M., Goldobin, D. S., Politi, A., and Torcini, A. (2021). in preparation

Shadlen, M. N. and Newsome, W. T. (1994). Noise, neural codes and cortical organization. Current opinion in neurobiology 4, 569–579

Shadlen, M. N. and Newsome, W. T. (1998). The variable discharge of cortical neurons: implications for connectivity, computation, and information coding. Journal of neuroscience 18, 3870–3896

Shibata, T. and Kaneko, K. (1998). Collective chaos. Physical review letters 81, 4116

Shu, Y., Hasenstaub, A., and McCormick, D. A. (2003). Turning on and off recurrent balanced cortical activity. Nature 423, 288–293

Smith, M. A. and Kohn, A. (2008). Spatial and temporal scales of neuronal correlation in primary visual cortex. Journal of Neuroscience 28, 12591–12603

Softky, W. R. and Koch, C. (1993). The highly irregular firing of cortical cells is inconsistent with temporal integration of random epsps. Journal of neuroscience 13, 334–350

Tiesinga, P. and Sejnowski, T. J. (2009). Cortical enlightenment: are attentional gamma oscillations driven by ing or ping? Neuron 63, 727–732

Ullner, E., Politi, A., and Torcini, A. (2018). Ubiquity of collective irregular dynamics in balanced networks of spiking neurons. Chaos: An Interdisciplinary Journal of Nonlinear Science 28, 081106

Ullner, E., Politi, A., and Torcini, A. (2020). Quantitative and qualitative analysis of asynchronous neural activity. Physical Review Research 2, 023103

van Vreeswijk, C. (1996). Partial synchronization in populations of pulse-coupled oscillators. Physical Review E 54, 5522

van Vreeswijk, C. and Sompolinsky, H. (1996). Chaos in neuronal networks with balanced excitatory and inhibitory activity. Science 274, 1724–1726. doi:10.1126/science.274.5293.1724

Volgushev, M., Chauvette, S., and Timofeev, I. (2011). Long-range correlation of the membrane potential in neocortical neurons during slow oscillation. Progress in brain research 193, 181–199

Whittington, M. A., Cunningham, M. O., LeBeau, F. E., Racca, C., and Traub, R. D. (2011). Multiple origins of the cortical gamma rhythm. Developmental neurobiology 71, 92–106

Whittington, M. A., Traub, R. D., Kopell, N., Ermentrout, B., and Buhl, E. H. (2000). Inhibition-based rhythms: experimental and mathematical observations on network dynamics. International journal of psychophysiology 38, 315–336

